# Predictive dendrites as a foundation for biological learning

**DOI:** 10.64898/2026.10.21.760638

**Authors:** Yuhang Song, Francesco Innocenti, Nima Mirkhani, Bariscan Bozkurt, Mara Ivan, Alif Aziz, Thomas Lukasiewicz, Andrew J. King, Claudia Clopath, Colin J. Akerman, Nicol S. Harper, Rafal Bogacz

## Abstract

A fundamental challenge for neuroscience is to identify laws of synaptic plasticity that are consistent with experimental observations at the level of individual neurons, whilst also explaining the ability of neural networks to learn complex tasks. Here we investigate a candidate law in which neurons adjust synaptic strengths to minimize the mismatch between somatic and dendritic activity. We demonstrate that models based on this rule can effectively adjust synaptic weights across network layers to learn non-linear tasks. Moreover, minimizing mismatch between spiking activity and dendritic inputs leads to integrate-and-fire dynamics and spike-timing-dependent plasticity. The model reproduces the experimentally observed dependencies of plasticity upon initial synaptic weights and spike patterns, whilst also generating a series of testable predictions. Our results show that a single principle – minimization of somato-dendritic mismatch – links learning at the network level with fundamental properties of individual neurons, including membrane potential dynamics and canonical features of synaptic plasticity.

---

Since the 1940s, our understanding of synaptic plasticity has been shaped by Hebb’s theory, which states that connections are strengthened between co-active neurons^1^. Hebbian plasticity has provided a framework for interpreting experimentally-observed phenomena such as long-term potentiation^2^ and spike-timing-dependent plasticity (STDP)^3^, whilst at the same time inspiring computational models of memory, such as Hopfield networks^4^. However, ‘classic’ Hebbian learning based on the co-activity of neurons^3,4^ is insufficient for many forms of learning. Traditional Hopfield networks have strikingly poor capacity for memorizing natural stimuli^5,6^ and training multi-layer networks for classification with Hebbian plasticity is ineffective^7^. These shortcomings of Hebbian plasticity result from problematic positive-feedback loops: As synaptic connections strengthen, neurons become increasingly co-active, triggering further potentiation that causes synaptic weights to grow towards their maximum values^8^. Hebbian plasticity also fails to account for several experimentally observed phenomena, including how synaptic plasticity can occur in the absence of postsynaptic spiking activity^9^, depends on the initial strength of a synapse^3,10^, and can vary depending on the location of the synapse^11^.

Within artificial neural networks, an effective learning algorithm known as backpropagation^12^ has been developed, which prescribes how the weights of synaptic connections should be modified to reduce the error at the network output. However, backpropagation is generally considered biologically unrealistic^13^ because synaptic weight changes depend on information originating from elsewhere in the network,whereas biological plasticity must rely on locally available signals. Although backpropagation can be approximated by predictive coding networks that rely on Hebbian plasticity^14^, these models require connectivity architectures that are inconsistent with cortical circuitry, including the absence of reciprocal excitatory connections.

A promising principle of biological learning is that synaptic plasticity acts to reduce mismatches between a neuron’s dendritic inputs and its somatic activity. We refer to this principle here as the predictive dendrites model (PDM), which builds upon related ideas previously explored in models where dendritic potentials predict a neuron’s somatic output^15^. Computational work has shown that networks containing predictive dendrites can learn effectively. Recurrent networks can function as associative memories and accurately retrieve large numbers of stored patterns^16^. Meanwhile, multi-layer networks can learn to classify images from machine-learning datasets, achieving performance levels competitive with backpropagation^17,18^. However, these models incorporated additional circuit mechanisms that either computed error signals analogous to those used in backpropagation^17^ or maximised the information encoded within each layer^18^. It therefore remains unclear whether minimization of somato-dendritic mismatch is itself sufficient to support effective learning without specialised circuit architectures or additional learning signals. Equally, it is unknown whether this same principle is consistent with fundamental properties of individual neurons and can account for experimentally observed features of synaptic plasticity. Resolving these questions is critical to establishing whether PDM could constitute a general biological principle of learning, capable of operating across diverse nervous systems and developmental stages.

Here we show that minimization of somato-dendritic mismatch is sufficient to support learning across multiple network layers, allowing networks of predictive neurons to learn non-linear associations without specialised error-computing circuitry. We then show that the same principle gives rise to integrate-and-fire dynamics in spiking neurons, linking PDM to fundamental properties of neuronal computation. Finally, we demonstrate that predictive dendrites reproduce experimental observations at the level of individual neurons and account for canonical features of synaptic plasticity, including phenomena that are difficult to explain using classical Hebbian principles. Together, these results establish somato-dendritic mismatch minimization as a candidate principle linking learning at the level of neural networks to the dynamics and plasticity of individual neurons.

## Results

### The predictive dendrites model

The PDM assumes that neurons seek to achieve consistency between their own somatic activity and the inputs they receive via different dendrites, which may come from different sources. The consistency between different dendritic inputs enables neurons to represent concepts that can be inferred from diverse cues (see example in Fig. 1a). In the PDM, individual dendrites can be thought of as predicting the somatic activity of a neuron. If there is a mismatch between dendritic potential and neural activity (Fig. 1b), the synaptic weights are modified to reduce the mismatch (Fig. 1c). For example, if a dendrite has a lower membrane potential than the soma, its connections from active inputs are strengthened, bringing the dendritic potential closer to the neural activity. The PDM still captures the general “fire together – wire together” principle of Hebb when connections between neurons are weak (Fig. 1d). However, critically, the weights are only modified when it is necessary for a dendritic prediction to match the neuron’s activity, so they may remain unchanged when neurons “fire together” but “are already wired” (Fig. 1d).

**Fig. 1.**
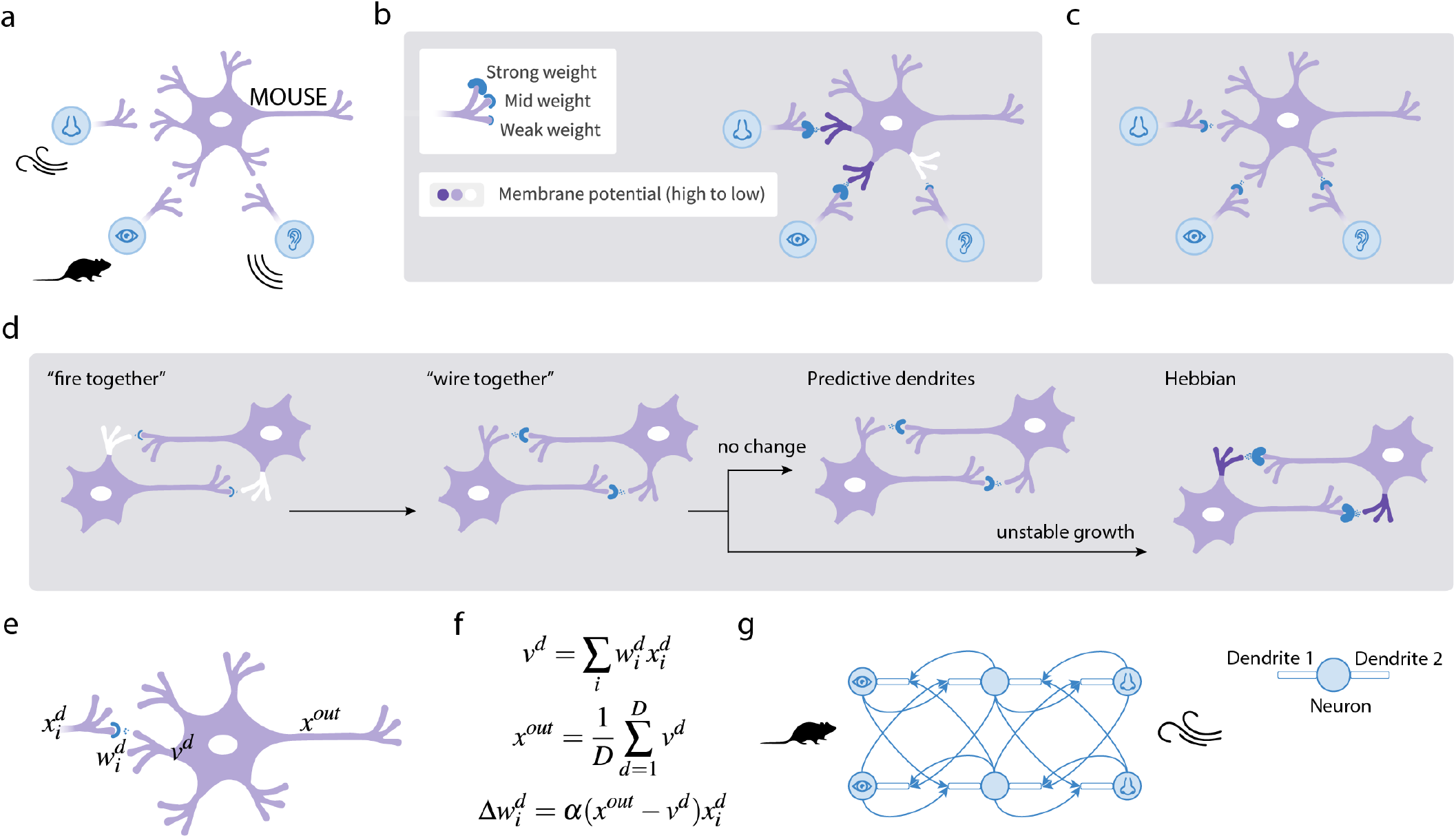
PDM. ▶ **a**| Example of a neuron that represents an abstract concept (a mouse), and which receives inputs from sensory areas encoding different modality cues associated with this concept via different dendrites. For the neuron to encode the concept, it should be similarly activated by each of these inputs, and the neuron’s activity should also be consistent with these inputs. ▶ **b** | Example of a situation where dendritic predictions do not match the activity of the neuron. ▶ **c** | Dendritic potentials in the neuron after the weights have been modified to reduce the dendritic errors. ▶ **d** | Relationship to Hebbian plasticity. The model increases the weights between co-active neurons, but only if dendritic errors are present. ▶ **e**| Notation used in the firing-rate formulation of the model: *v*^*d*^ denotes the membrane potential in dendrite 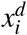 is the activity of input neuron *i* projecting to dendrite *d*, with 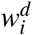 being the weight for this input, and *x*^*out*^ is the output or activity of the neuron. ▶ **f** | Firing-rate formulation of the linear PDM, where *D* is the number of dendrites. ▶ **g**| Network architecture. Circles denote neurons, rectangles denote dendrites, and arrows denote connections between the neurons.

To analyse learning in the PDM, we first consider a simple formulation, where we define the *activity* of a neuron as its firing rate. The linear version of this model can be described by only three equations (Fig. 1e-f). The first equation states that the membrane potential of a dendrite is the sum of its inputs scaled by synaptic weights. According to the second equation, the activity of a neuron is determined by the average of the membrane potentials of its dendrites. Finally, the third equation states that the change in a synaptic weight on a particular dendrite is a product of: (i) the dendritic error, defined as the difference between the neuron’s activity and the dendritic potential; (ii) the activity of a presynaptic neuron; and (iii) a learning rate parameter (*α*).

In Methods Section A we show that the equations for somatic activity and synaptic weight update in Fig. 1f arise from minimization of the sum of squared dendritic errors, which we refer to as a neuron’s ‘energy’ (the word ‘energy’ is used to describe a function that neural systems minimize^4,19^ by analogy to potential energy minimised by physical systems). Specifically, the activity equal to the average of dendritic potentials corresponds to the activity minimising the energy, and the weight update most directly reduces the energy (i.e., is proportional to its negative gradient). Such optimization of an energy by both activity and plasticity is a feature of many biologically plausible ‘energy-based models’^19^, which are known to learn well^20^.

The neurons can be connected into networks that learn associations between stimuli. In such networks, different sensory stimuli – in this case from different modalities – can be encoded by neurons at the two ends of the network (Fig. 1g). Throughout this paper we consider networks in which neurons are organized into layers. When the network predicts one modality from another, the neurons on the left of Fig. 1g can be thought of as the input to the network, the neurons on the right as the output, and the neurons in between are referred to as hidden. We consider a simple case where the hidden neurons have just 2 dendrites receiving inputs from the populations encoding the 2 modalities, while the input and output neurons have only a single dendrite as we assume that sensory stimuli can directly determine the activity of these neurons. When the network is being trained to produce a particular target output for a given input, both input and output neurons are fixed to the training patterns, the hidden neurons settle their activity, and the weights are updated for all neurons (details in Methods Section B).

### Predictive dendrites enable learning in networks with multiple layers

The PDM can effectively modify the weights across multiple layers to reduce the error at the output. This capacity arises because the network is recurrently connected; when a target output is provided, it changes the activity in multiple layers (Fig. 2a-c). The resulting activity pattern in the network is then reinforced by plasticity reducing dendritic errors (Fig. 2d, further intuition provided in Supplementary Fig. 1). In Supplementary Information Sections 1-2, we demonstrate analytically that a linear PDM modifies weights in all layers, such that the output of the network changes towards the target of training (under certain conditions). Moreover, in Supplementary Information Section 3, we show that the weight modification in the model under certain conditions – corresponding to preserving representation of inputs in hidden layers – approaches that in bidirectional predictive coding, which has been shown to achieve backpropagation-level accuracy^21^.

**Fig. 2.**
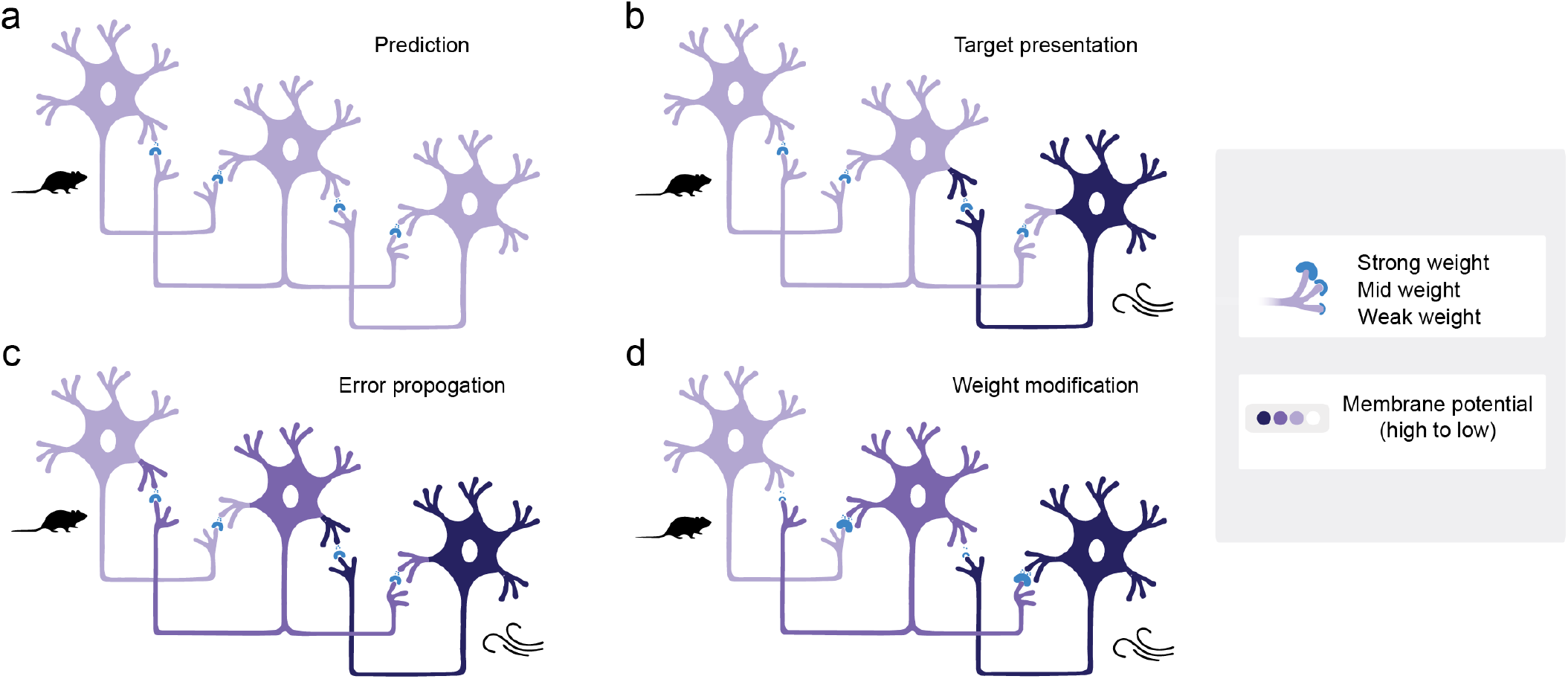
Error propagation in a network of neurons with predictive dendrites. Each panel schematically illustrates the activity in a simple network of three interconnected regions or layers with one neuron shown per layer. The left neuron is located in a visual region, the right in an olfactory region, and the middle in an association layer connecting the two sensory regions. The activity in different parts of neurons is denoted by colour and the strengths of synaptic connections by the sizes of blue semi-circles (see key). ▶ **a**| When a visual input is presented, it activates a visual neuron (left), and this activity propagates through existing connections to other neurons, generating a prediction for the activity expected in the olfactory layer. ▶ **b** | When an olfactory stimulus arrives, it triggers a different activity than that predicted, and the network needs to learn to predict the target activity driven by the stimulus. ▶ **c**| The target activity in the olfactory region changes the activity in the middle association layer, resulting in the propagation of error, since dendritic prediction errors arise in all layers (shown by difference in colours of dendrites and adjacent soma). ▶ **d** |Weights are modified to reduce dendritic errors, resulting in stronger weights from visual to olfactory layers. If the resulting network were to predict the olfactory stimulus from visual one, it would produce higher activity of the olfactory neuron, i.e., closer to the target.

To numerically confirm that the PDM can indeed propagate errors and appropriately adjust weights in multiple layers, we trained it on non-linear problems known to require networks with at least one hidden layer, starting with the classic XOR problem (Fig. 3a). To solve such tasks, the PDM needs to be extended to include a non-linearity. Therefore, we assume that the activity of neurons cannot be negative, and set it to 0 if the average of the dendritic membrane potentials is negative (equivalent to the “ReLU” activation function; Fig. 3b, middle equation). For non-linear neurons, minimization of the corresponding augmented energy yields an additional gate on plasticity: weights of a hidden unit are modified only when its activity is positive (Fig. 3b, bottom equation; Supplementary Information Section 4). This enables neurons to learn to represent useful features of stimuli, which requires them to update their weights to make their response closer to the input only when the feature is present (Methods Section A). The weight update equation in Fig. 3b can be viewed as an extension of Hebbian learning, as the weight change depends on the activity of the presynaptic neuron, the activity of the postsynaptic neuron, and the prediction error of the dendrite. The non-linearity can be introduced into the energy function in other ways (example given in Supplementary Fig. 2), but we focus on the formulation resulting in weight update in Fig. 3b because its generalization to spiking neurons will produce STDP.

**Fig. 3.**
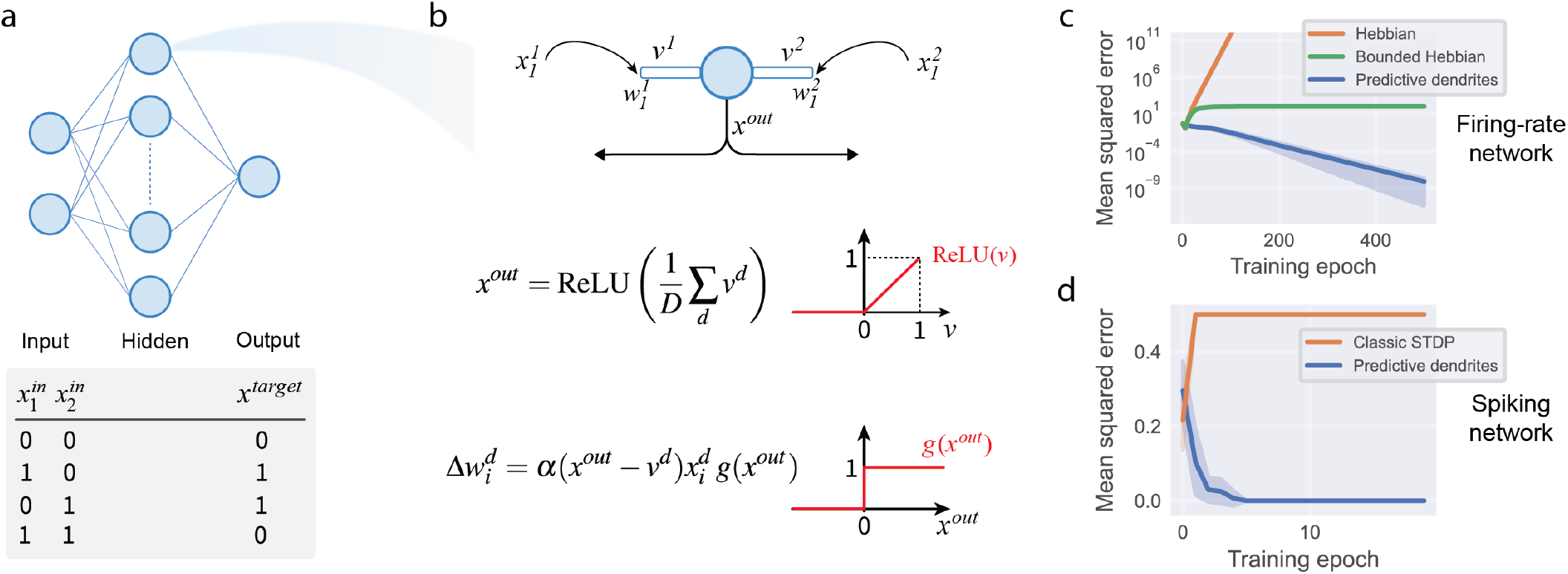
Learning the XOR problem. ▶ **a**| Network and training set for the XOR problem. ▶ **b**| Detail of a single hidden unit. Diagram at the top summarizes the notation, and the equations below describe activity and weight update for the non-linear model. The weight-update equation includes the additional term *g*(*x*^*out*^), which is equal to 1 only when the neuron has positive activity, as shown in the bottom graph. ▶ **c** | Training loss of the firing-rate PDM and Hebbian baselines during learning. ▶ **d**| Training loss of the spiking PDM and a classic online STDP model^22^ during learning. The training error of the PDM reaches zero because it is defined as a squared difference between the produced and desired number of spikes (Methods Section J), and therefore takes only discrete values.

The PDM with one hidden layer was able to learn the XOR task, as its training error decreased to a low level during learning (Fig. 3c). By contrast, models based on standard Hebbian learning (Methods Section D) could not solve the task (Fig. 3c). If a network has a sufficient number of non-linear hidden neurons, the XOR problem can be solved by simply learning the hidden-to-output weights, but the PDM additionally adjusted the input-to-hidden weights such that the hidden neurons represent features required to solve the task (Supplementary Fig. 3a-c).

While training a basic version of the PDM in other tasks, we found that the weights of hidden neurons could converge to two trivial configurations, which reduced their local energy but did not result in the network making correct predictions. Specifically, the weights feeding into a hidden neuron can converge to 0, and all the hidden neurons may encode the same information. These problems are forms of “representation collapse”, a common failure mode in self-supervised learning models, and preventing such collapse typically requires additional mechanisms^23–25^. In Supplementary Fig. 4 we analyse two simple tasks in which these trivial configurations occur and show that they can be prevented by biologically plausible mechanisms, including activity regularization and lateral connectivity within a layer, both of which are supported by multiple circuit and cellular mechanisms in the brain^26–28^. We have also demonstrated the capacity of a variant of the PDM to learn a standard machine learning benchmark involving classification of species of iris (Supplementary Fig. 5).

Benchmark tasks typically used to assess learning abilities of computational models involve classification of images. Although previous work has demonstrated that these tasks can be learned by models based on predictive dendrites principle with additional circuit mechanisms^17,18^, we found that the standard PDM is unable to learn these tasks (Fig. 4a), even with the extensions mentioned above. One element included in a previous model^18^ was a contrastive learning rule, which requires an additional weight modification based on activity arising when only input is provided (Fig. 4b). Although this additional “free phase” reduces the simplicity and temporal locality of the original learning rule, it is useful to consider because the free-phase contribution vanishes in closely related models^29^, in which there are no predictions errors if only input is provided. Dendritic errors also vanish in the PDM when only input is constrained in the examples shown in Fig. 1 and Supplementary Fig. 1. Consequently, it is useful to analyse contrastive learning in the PDM, because if future work identifies conditions under which the free-phase contribution vanishes, there is a potential to recover a single-phase rule without sacrificing performance. Here we asked whether a PDM with a simpler architecture (Fig. 1g) could learn these tasks using a contrastive version of the rule, and found it also approaches performance similar to backpropagation (Fig. 4a, Methods Section C). The model outperformed a network with fixed input weights (Fig. 4a) demonstrating that it also refined the representation in the hidden layer (see Supplementary Fig. 6 for additional models and ablations).

**Fig. 4.**
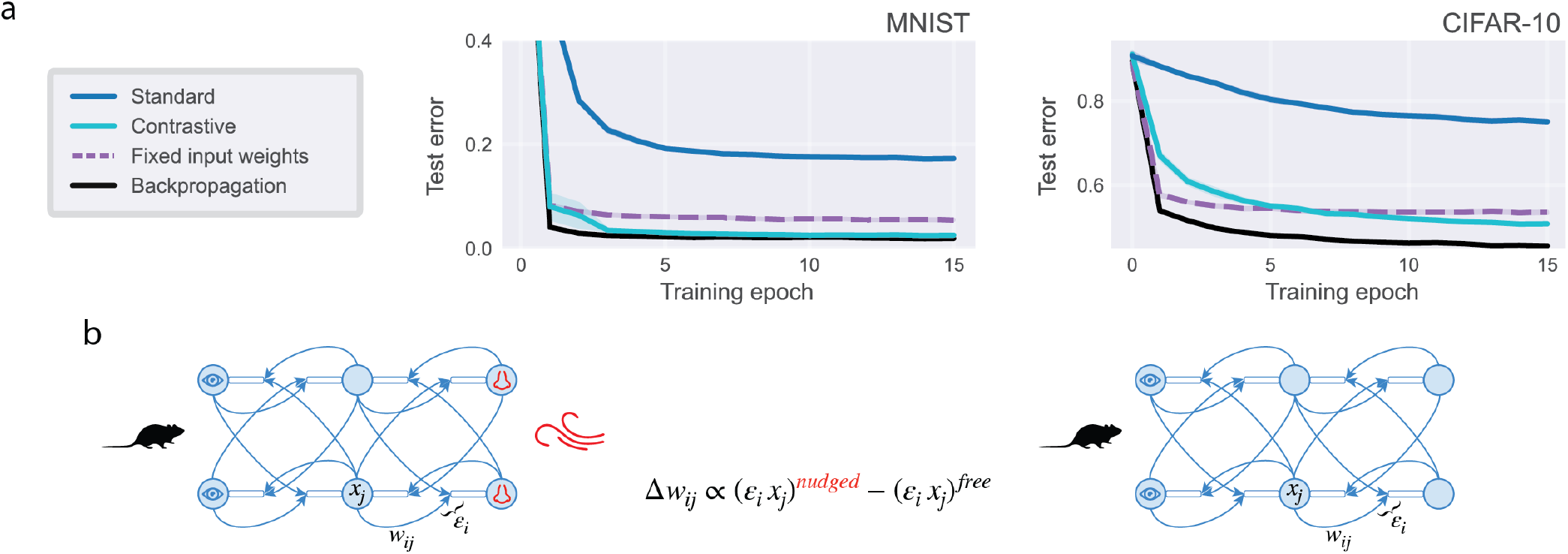
Performance of the contrastive variant of the PDM in classification of benchmark datasets. ▶ **a**| Classification test error on MNIST (handwritten digits) and CIFAR-10 (images of common animals and vehicles) datasets of a variant of the PDM trained with a contrastive learning rule, compared to the standard model and two backpropagation baselines (Methods Section E). ▶ **b**| Schematic illustration of the contrastive rule. As described in detail in Methods Section C, the contrastive variant of the PDM modifies weights by contrasting updates under two different relaxed states of the network: (i) a “free phase” where the output of the network is unclamped and left free to vary; and (ii) a “nudged phase” where the dynamics are nudged toward, or slightly influenced by, the target.

In summary, the PDM can update the weights across layers and provides a foundation on which other features that improve performance can be built. We next consider whether the PDM can capture experimental data on neuronal dynamics and STDP.

### Minimization of dendritic prediction errors results in integrate-and-fire dynamics

We considered the possibility that if PDM represents a fundamental biological learning principle, key properties of neurons should follow naturally from its underlying assumptions. A fundamental feature of the dynamics of biological neurons is integration of synaptic inputs in the membrane potential until reaching a spiking threshold. Building on earlier work^30^, here, we show that the integration to threshold naturally arises if a neuron produces spikes in a way that minimizes dendritic prediction errors. The resulting spiking model formulation is summarised and illustrated with a simple simulation here, while the mathematical details are given in Methods Section F.

We consider a simple case where the neuron contains just two dendrites, each receiving input from a single neuron, and Fig. 5a shows variables describing different elements of the neuron. Fig. 5b shows the values of model variables in a sample simulation. In the spiking formulation, the neurons produce spikes, and we denote the spikes on input neurons projecting to dendrite *d* by *s*^*d*^ (Fig. 5b, Input spikes). Despite spikes being produced in discrete intervals, their effect on the postsynaptic neurons lasts longer, as they produce postsynaptic potentials. We therefore define the input signals *x*^*d*^ as the temporally extended effect of spikes. Hence, the signal increases whenever the spike is produced, and then decreases exponentially with a decay rate determined by a parameter *λ* (Fig. 5b, Input signal). The membrane potentials within dendrites *v*^*d*^ are determined by the input signals scaled by the associated weights, and delayed by a parameter *τ*, which corresponds to a synaptic transmission delay (Fig. 5b, Dendritic potential).

**Fig. 5.**
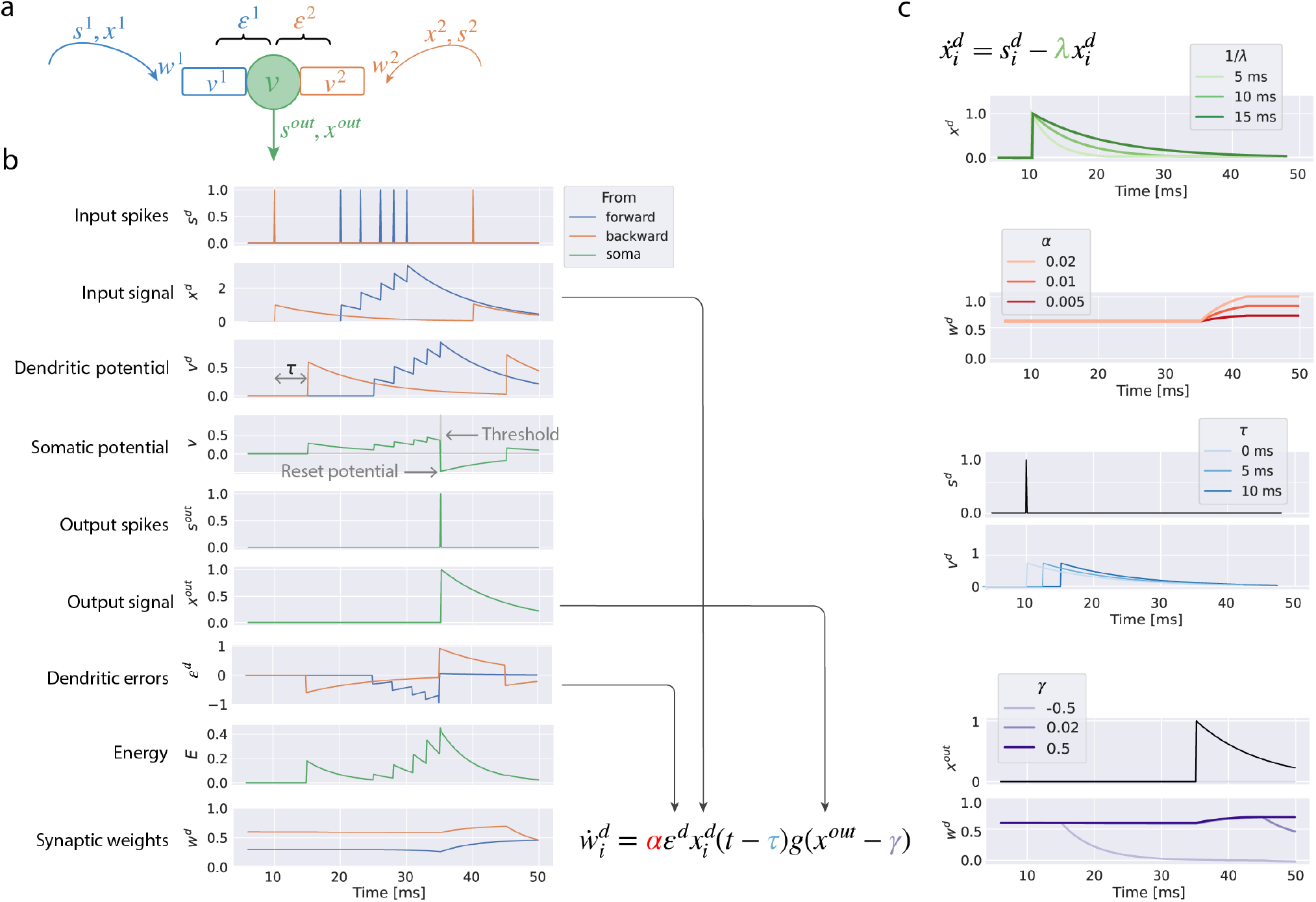
Spiking formulation of the PDM. ▶ **a** | Summary of variables describing different components of a neuron. ▶ **b**| Evolution of the variables during a simulation. All parameters of the neuron were set to default values (Methods Section F), except for the learning rate which was increased to *α* = 0.1 to make weight changes more visible. The variables associated with the two dendrites are shown in blue and orange, while the variables associated with the soma and axon are in green. In the equation describing the weight-update rule, arrows indicate the variables involved. ▶ **c**| Effects of parameters on model dynamics. The panel visualizes simulations similar to panel b where each display varies a particular parameter, and the graphs show the variable(s) most directly affected by the parameter (for simplicity, we only show variables associated with the first dendrite). Parameter *λ* determines how rapidly membrane potential decays. Parameter *α* affects how much weights change during learning. Parameter *τ* denotes the delay between the presynaptic spike and a change in dendritic potential. Parameter *γ* denotes the minimal level of output signal required for plasticity, hence its sign determines whether postsynaptic activity is necessary for plasticity.

We now consider how a postsynaptic neuron should behave to minimize dendritic errors. These errors *ε*^*d*^ = *x*^*out*^− *v*^*d*^ are defined as the difference between a neuron’s output signal *x*^*out*^ and the dendritic potential *v*^*d*^, where the output signal is defined analogously to the input signal (i.e. it increases with spikes and decays afterwards). The output signal plays a role equivalent to a neuron’s activity in the firing-rate formulation. However, a spiking neuron cannot control its output signal directly, but only “decide” whether to spike or not. Hence, we assume, as in previous work^30,31^, that a neuron produces a spike if and only if the spike reduces the energy (defined as a sum of squared dendritic errors). Such spikes should be produced when the dendritic potentials become so large that prediction errors can be reduced by increasing neuron’s output. In Methods Section F, we show that the spikes should be produced when the difference between the average dendritic potentials and the neuron’s output reaches a threshold, and we denote this difference by 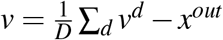. We identify *v* with the somatic membrane potential because it has the characteristic dynamics of a membrane potential: synaptic inputs increase *v* until it reaches a spike threshold^31^ (Fig. 5b, Somatic potential). Moreover, *v* rapidly decreases after a spike^31^ because when the spike *s*^*out*^ is produced (Fig. 5b, Output spikes), the output of the postsynaptic neuron *x*^*out*^ increases (Fig. 5b, Output signal). The spike also reduces the absolute prediction error associated with the more activated dendrite (Fig. 5b, Dendritic errors) and the energy (Fig. 5b, Energy). In summary, the resulting dynamics correspond to an integrate-and-fire model^32^, which emerges here from minimization of dendritic errors.

The synaptic plasticity of the spiking model is analogous to the firing-rate version (equation in Fig. 5b). The weight updates on a dendrite *d* are determined by the dendritic error *ε*^*d*^ multiplied by the presynaptic activity *x*^*d*^ and the learning rate parameter *α*. Similar to the non-linear firing-rate model, the weights are only modified if the postsynaptic neuron has sufficient output signal. Since after a spike *x*^*out*^ never reaches 0 (as it decays exponentially), we introduce a parameter *γ* describing the minimum output signal required for plasticity. Consequently, the weights remain constant until the postsynaptic neuron produces a spike (Fig. 5b, Synaptic weights). After the spike, both weights increase because there are positive prediction errors associated with both dendrites. When the potential in the second dendrite *v*^*d*^ increases at *t* = 45*ms*, the weight *w*^2^ decreases due to a negative prediction error for the second dendrite. The spiking formulation has four parameters, and Fig. 5c illustrates the effect of each parameter on model behaviour. We now turn to investigating if the modification of synaptic weight, which minimizes somato-dendritic mismatch, can capture canonical properties of synaptic plasticity.

### The PDM reproduces spike-timing-dependent plasticity

Since the learning rule of the spiking model (Fig. 5b) depends on temporally filtered pre- and postsynaptic signals, its weight changes depend naturally on relative spike timing. Here, we demonstrate that when the model is simulated in a paradigm employed to study STDP, it can reproduce experimentally observed shapes of STDP curves.

We applied an STDP experimental protocol, where pre- and postsynaptic neurons are stimulated, to a simple model of a neuron with two dendrites, considered in the previous section (Fig. 5a). The weights to the neuron had low values (*w*^*d*^ = 0.5) so that a single presynaptic spike did not trigger a postsynaptic spike. We provided a single presynaptic spike as input to the first dendrite, which was followed or led by a single spike we triggered in the postsynaptic neuron. We then plotted the percentage change in the weight *w*^1^ as a function of the delay between these two spikes.

The model produced the canonical STDP profile (Fig. 6b). To gain insight into why it produced this curve, Fig. 6a and Fig. 6c show the evolution of the key model variables for the presynaptic-postsynaptic (pre-post) delays that gave the most pronounced decrease and increase in the weight, respectively.

**Fig. 6.**
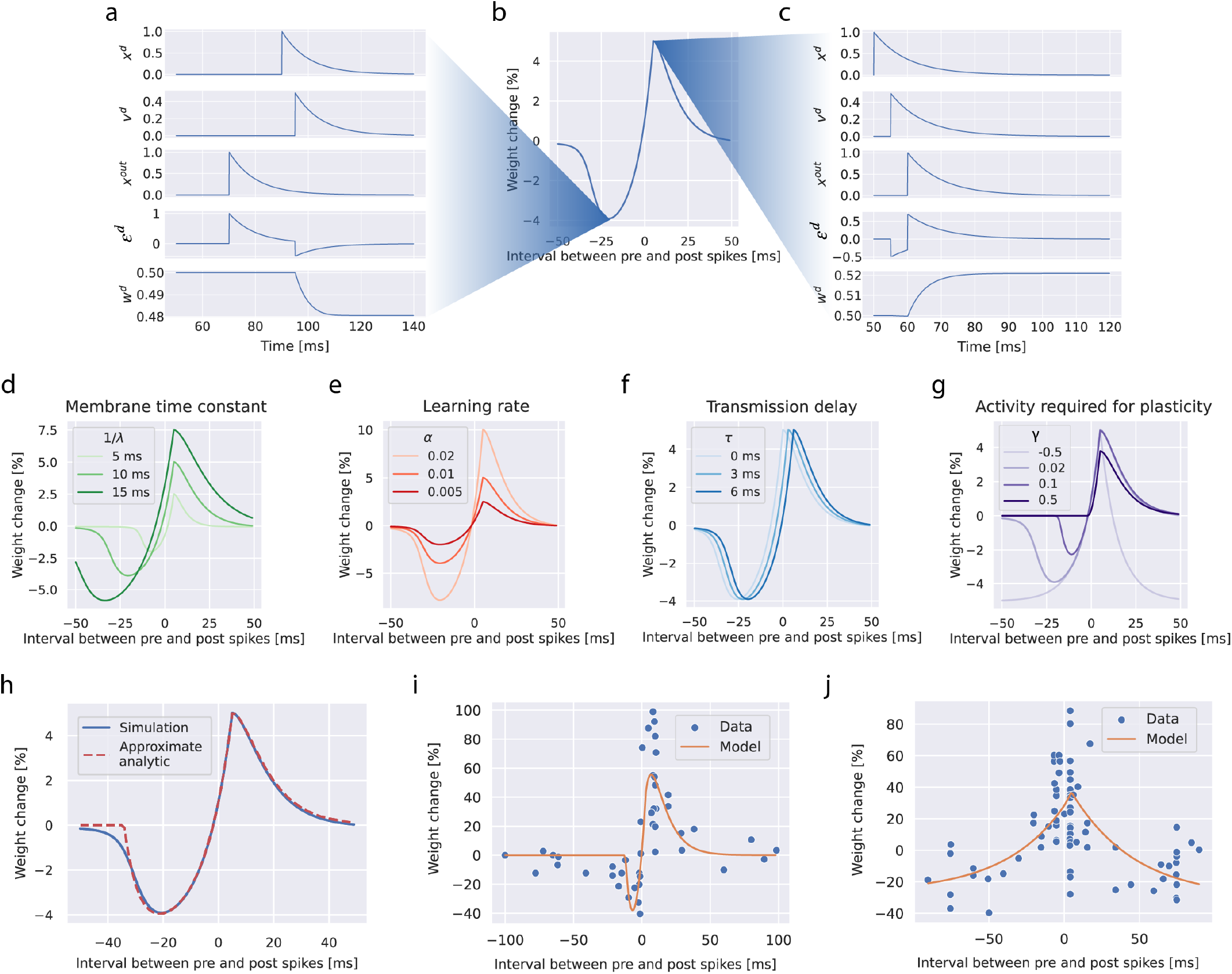
The PDM reproduces STDP curves. ▶ **a-c** | An STDP curve produced for default model parameters (b), and the changes in model variables when the pre-post delay is −20ms (a) and 10ms (c). ▶ **d** | The decay rate (*λ*), which determines the membrane time constant 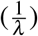, scales both the height and the width of the curve. ▶ **e** | The learning rate scales the weight changes. ▶ **f** | The synaptic transmission delay shifts the curve. ▶ **g** | The effects of the minimum postsynaptic activity required for plasticity (*γ*) depend on its value. Negative *γ* results in plasticity that does not require postsynaptic activity and in a symmetric STDP curve. Conversely, a high value (*γ* = 0.5) means that plasticity can only occur just after a spike when *x*^*out*^ is high and prediction error is positive; thus, the STDP curve only contains a positive part. Positive *γ* values near zero regulate how deep and wide the negative part of the STDP curve is. ▶ **h**| Comparison of a simulated STDP curve with an analytical approximation. ▶ **i-j**| Comparison of experimental data on STDP (from^33^ and^34^ respectively) with best-fitting curves from the PDM.

According to the learning rule in Fig. 5b, the synaptic weight is modified only after both presynaptic and postsynaptic spikes occur (when both *x*^*d*^(*t*− *τ*) > 0 and *g*(*x*^*out*^ − *γ*) > 0). This condition was met from 95ms and 60ms in Fig. 6a and Fig. 6c, respectively. When these conditions are met, the direction of weight change depends on the sign of dendritic error, which was negative and positive in Fig. 6a and Fig. 6c, respectively.

In the PDM, the shape of the STDP curve is determined by the values of model parameters, and Fig. 6d-g illustrate the effect of changing each of the 4 parameters. In Methods Section G, we derive an analytic formula for the STDP curve that closely approximates the simulation (Fig. 6h), and in which the parameter effects are directly apparent.

We next investigated whether the model can quantitatively capture experimentally measured STDP curves. This is not *a priori* guaranteed, since unlike previous models^35,36^, the PDM was not designed to capture STDP curves, but rather the underlying computations and learning process of neurons (Fig. 5b). Nevertheless, after optimizing its parameters, the model quantitatively fitted the experimentally measured STDP curves (Fig. 6i-j, Methods Section H).

The spiking version of the PDM retains the ability to learn non-linear problems and modify weights throughout the network to minimize the output error. As an example, we trained networks with one hidden layer on the XOR problem (Methods Section I). The network learned the task rapidly (Fig. 3d). The network also learned features required to solve the XOR problem in the hidden layer (Supplementary Fig. 3d). By contrast, a network with plasticity described by the classic STDP model^22^ (Methods Section J) was unable to learn the task, because certain weights diverged to their maximum values, as they were increased whenever the postsynaptic spike followed the presynaptic spike. Although both models compared in Fig. 3d reproduced STDP, only in the PDM did the weight modification depend on the dendritic errors, which enabled the model to learn the task.

### The PDM captures the effects of synapse state and spike patterns upon synaptic plasticity

The PDM also captures a well-established observation that long-term potentiation is easier to produce for synapses that are initially weak, rather than strong^3,10,37,38^. This effect naturally emerges from the model because a higher initial weight results in a larger dendritic prediction, and hence a smaller prediction error and weight change. Fig. 7a illustrates the effect by showing how the STDP curve from the PDM depends on the initial weight, and that the peak of the curve is lower for higher initial weights. Fig. 7a also shows that the trough of the curve is slightly more pronounced for high initial weights. This occurs because a high initial weight produces a high prediction and thus more negative dendritic error after a presynaptic spike. Nevertheless, the effect on the trough is much smaller than on the peak, because the y-axis in Fig. 7a corresponds to the percentage of weight change (Δ*w/w*_*initial*_), and the larger absolute weight reduction for higher initial weights is scaled down by normalization to the initial weight.

**Fig. 7.**
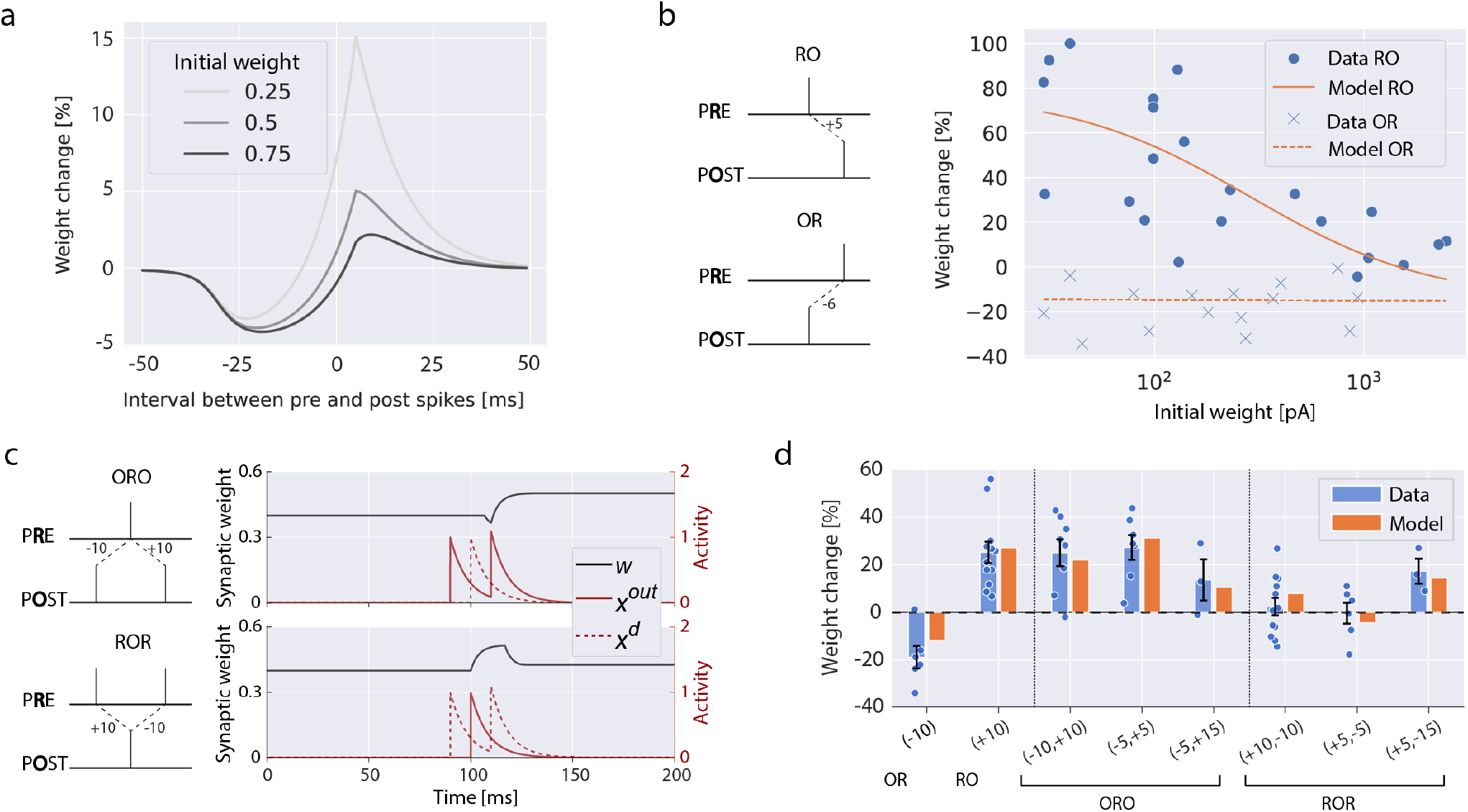
Dependence of plasticity on initial synaptic strength and spike pattern. ▶ **a** | Effect of the initial weight on the shape of STDP curve. ▶ **b** | Comparison with experimental data on the dependence of weight change on the size of initial synaptic response (from^3^) with best-fitting curves from the PDM. The filled circles show data from the condition in which the presynaptic neuron was stimulated first, and the postsynaptic neuron was stimulated after a delay of 5 ms (illustrated in top-left diagram). The crosses show data from the condition with the opposite order, i.e. in which the postsynaptic neuron was stimulated first, and the presynaptic neuron was stimulated after a delay of 6 ms (bottom-left diagram). ▶ **c** | Weight changes under spike triplet paradigms: post-pre-post (top) and pre-post-pre (bottom). ▶ **d** |Comparison of experimental data for single-pair and triplet conditions from^39^ (mean ± SEM, individual datapoints shown) with weight changes in the fitted PDM. Each condition is labelled by its spike timing interval(s) in ms, where positive and negative values indicate pre-post and post-pre pairings, respectively.

Fig. 7b shows that the PDM with fitted parameters reproduced the experimentally observed effects of initial synaptic strengths on plasticity^3^ (Methods Section H). In particular, the model captured both the reduced potentiation of synapses with higher initial weights and the negligible effect of initial weights on depression^3^.

While classic STDP protocols characterize plasticity using repeated single pre-post spike pairings, neurons in vivo are subject to complex spike trains that may simultaneously engage both potentiation and depression processes^33,39^. The simplest paradigm to probe such interactions uses triplets of spikes, which comprise pre-post-pre or post-pre-post combinations, for example. The plasticity induced by these triplets has features that cannot be explained by basic STDP alone, and have been shown to require specific modifications to STDP models^40,41^. By contrast, the spiking PDM requires no change to the learning rule in order to handle arbitrary spike patterns, allowing pair and triplet effects to arise from the same rule. Under equal pre-post and post-pre intervals in triplets, PDM produced qualitatively different outcomes depending on the temporal order of the pairs (Fig. 7c). Specifically, in the post-pre-post condition, the depression due to the initial post-pre pair started a few milliseconds after the presynaptic spike because of a synaptic transmission delay, and was rapidly interrupted by the increase due to the subsequent pre-post pair, resulting in an overall weight increase. Meanwhile, in the pre-post-pre condition, the increase and decrease in weights cancelled one another, resulting in little change overall. The PDM reproduced the experimentally observed asymmetric integration of potentiation and depression across various triplet conditions (Fig. 7d)^39^. In summary, we demonstrated that the PDM can account for the plasticity due to diverse stimulation patterns.

## Discussion

Our findings reveal that a single principle – minimization of somato-dendritic activity mismatch – can link learning in networks to canonical properties of individual neurons, including integrate-and-fire dynamics and several experimentally observed features of synaptic plasticity. The results therefore demonstrate that phenomena often described separately at the network, cellular and synaptic levels, can in fact arise from a common learning objective.

### Relationship to experimental data

We have shown that experimentally observed STDP curves emerge when the spiking PDM is simulated in classical plasticity-inducing protocols. These results support the view that STDP is not a basic law of synaptic plasticity, but rather a consequence of more fundamental learning rules^41^. By extension, the PDM can also account for other experimentally observed features of synaptic plasticity. For instance, it has been observed that the magnitude of synaptic potentiation tends to be greater when the postsynaptic neuron produces a burst of multiple spikes, rather than a single spike^42^. This fully complies with the PDM, because multiple spikes correspond to larger postsynaptic activity, which would produce a larger dendritic error and hence a stronger weight increase. In addition, experiments have revealed that activating a presynaptic neuron when the postsynaptic neuron remains inactive tend to lead to long-term depression^9,38,43^. The PDM can reproduce this pattern (if parametrized such that plasticity is not gated by postsynaptic activity, *γ* < 0), because activating inputs to a dendrite without subsequent postsynaptic activity produces a negative dendritic error. Activating a presynaptic neuron followed by a postsynaptic neuron has been shown to strengthen synapses close to the soma of layer 5 pyramidal neurons in cortex, but weaken synapses farther from the soma^9^. One explanation for this effect within the PDM is that the output signal, *x*^*out*^, is conveyed to synapses by backpropagating action potentials, whose amplitude attenuates with dendritic distance^44^. Hence, *x*^*out*^ that is read out by a synapse close to the soma may be large, resulting in a positive dendritic error, whereas *x*^*out*^ read out far from the soma may be small, yielding a negative dendritic error.

A support for the key assumption of the PDM, that prediction errors are encoded in somato-dendritic mismatch, has been provided by a recent study which demonstrated that the difference between somatic and dendritic activity accurately tracks behavioural errors made by mice, and that perturbing this difference results in disrupted learning^45^.

### Relationship to other models

Among many theories describing learning in hierarchical cortical networks^14,46–50^, here we focus on discussing work most related to the PDM. A family of theories assumes that prediction errors are encoded in the activity of apical dendrites, which convey top-down information^17,51–54^ (rather than their mismatch with soma). We do not view these theories conflicting with the PDM, because we view the PDM as a description of the properties of more conventional dendrites, such as the basal dendrites of pyramidal neurons, which receive most of the feedforward afferent input. Since apical dendrites have unique properties^55^, they may indeed support more sophisticated mechanisms built on the foundation of the PDM. The PDM is related to predictive coding networks^19,56^ and the symmetric predictive estimator^57^.

These models also compute prediction errors, but in the PDM these errors are computed within a neuron, and directly regulate its plasticity. This removes unrealistic constraints on connectivity present in predictive coding networks. For example, in these networks no two neurons are reciprocally connected via excitatory connections^56^, as they would grow unstably under Hebbian plasticity, while reciprocal excitatory connections between neocortical neurons have been commonly observed experimentally^58,59^. The spiking PDM is related to several previous models of STDP. It shares an important feature with the voltage-based STDP model^36^, in that that plasticity only occurs if the postsynaptic neuron has a minimal level of activity. It has previously been demonstrated that STDP can arise in other models minimizing prediction errors^60,61^. These models differ from the PDM in the definition of prediction errors^60^ or factors determining the shape of the STDP curve^61^, and could be distinguished in experimental approaches suggested below.

### Experimental predictions and future directions

The PDM makes several unique experimental predictions summarized in Fig. 8. If future experiments confirm that the principles of the PDM describe biological synaptic plasticity, then subsequent work could investigate the mechanisms through which the dendritic errors are computed within a cell. It would also be interesting to study the scale of the neural components generating predictions and prediction errors, i.e. whether they correspond to individual spines, dendritic branches, entire dendrites, or even groups of dendrites, and whether behavioural and *in vivo* physiological measures of learning can be accounted for in terms of the PDM.

**Fig. 8.**
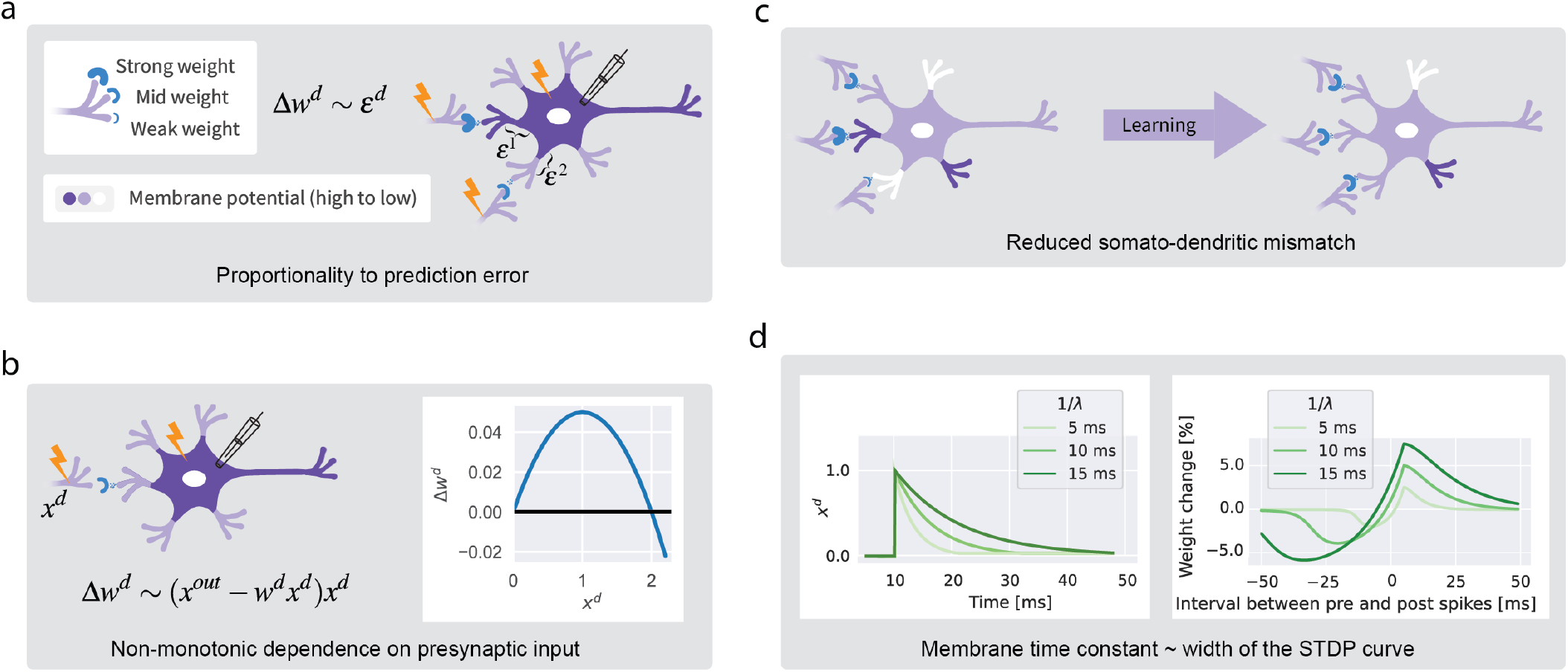
Experimental predictions. ▶ **a**| The PDM predicts that the synaptic modification depends on dendritic error. This prediction could be tested by varying inputs to a neuron and its somatic activity, and observing the effects of dendritic errors on synaptic weight modification. ▶ **b**| The model predicts that the change in synaptic weights to a dendrite should depend non-monotonically on the presynaptic input to the dendrite. To visualize this, consider a simple case of a dendrite receiving a single input. The change in synaptic weight associated with this input (Fig. 1f) then becomes Δ*w*^*d*^ = *α*(*x*^*out*^ − *w*^*d*^*x*^*d*^)*x*^*d*^. This is a quadratic function of *x*^*d*^, shown on the right for sample parameters (*x*^*out*^ = 1, *w*^*d*^ = 1 and *α* = 0.1). This dependence of plasticity on presynaptic input could be tested by varying the input *x*^*d*^ across experimental conditions (while keeping the postsynaptic activity *x*^*out*^ fixed at a non-zero value). The model predicts no plasticity without input, and as the input is increased across conditions, the weight change Δ*w*^*d*^ should initially increase across conditions, then reach a maximum, and even decrease (due to a negative dendriticerror). ▶ **c** | The model predicts that the somato-dendritic mismatch should reduce for dendrites receiving inputs representing learned stimuli. This prediction could be tested by imaging dendrites and soma during a learning task. ▶ **d**| The model predicts that the shape of the STDP curve depends on parameters (Fig. 6d-g) that also determine other properties of the neuron (Fig. 5c). For example, *λ* describes the decay rate of the postsynaptic potentials (Fig. 5c replotted on the left) due to the leak through the neuron’s membrane^62^; hence 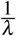 corresponds to the time constant of the membrane potential. The model predicts that this parameter also determines the width of the STDP curve (Fig. 6d replotted on the right); hence the width of the STDP curve should be proportional to the membrane time constant of the postsynaptic neuron.

## Acknowledgements

Presented research was supported by the following grants: Wellcome Trust grant 313955/Z/24/Z (RB, CA, CC, AK), China Scholarship Council under the State Scholarship Fund (YS), J.P. Morgan AI Research Awards (YS), Biotechnology and Biological Sciences Research Council grant BB/S006338/1 (RB), Medical Research Council grants MC UU 00003/1 and UKRI936 (RB), the Alan Turing Institute under the EPSRC grant EP/N510129/1 (TL), the AXA Research Fund (TL), Wellcome Trust Principal Research Fellowship WT108369/Z/2015/Z (AK, NH). Some of the simulations were supported by Google Cloud Credits awarded to F.I. by the Google TPU Research Cloud Programme (2026). The funders had no role in study design, data collection and analysis, decision to publish or preparation of the manuscript.

## JPMORGAN CHASE & CO

This research was funded in part by JPMorgan Chase & Co. Any views or opinions expressed herein are solely those of the authors listed, and may differ from the views and opinions expressed by JPMorgan Chase & Co. or its affiliates. This material is not a product of the Research Department of J.P. Morgan Securities LLC. This material should not be construed as an individual recommendation for any particular client and is not intended as a recommendation of particular securities, financial instruments or strategies for a particular client. This material does not constitute a solicitation or offer in any jurisdiction.

## Author contributions

Y.S. and R.B. conceived the project. Y.S., F.I., N.M., B.B., M.I., A.A., N.H. and R.B. performed simulations. Y.S. and R.B. performed mathematical analyses. Y.S., T.L., N.H. and R.B. provided supervision. Y.S., F.I., N.M., B.B., M.I, and R.B. wrote the paper. A.K., C.C., C.J.A. and N.H. revised the paper.

## Methods

### A Firing-rate formulation for individual neurons

We will first derive here the linear model summarised in Fig. 1f, and then show how the model can be extended to include a non-linearity. Using the notation introduced in Fig. 1e, we defined the membrane potential in dendrite *d* as (the first equation in Fig. 1f, rewritten here for convenience):

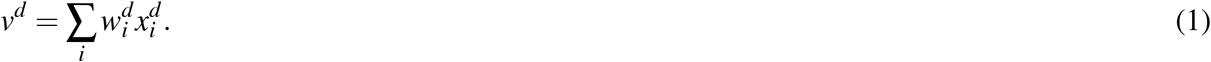

We can define an energy function of a neuron as a sum of squared dendritic prediction errors over all *D* dendrites of the neuron:

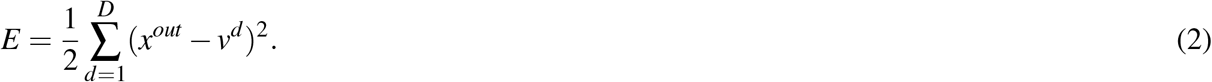

Assuming that the neuron changes its activity *x*^*out*^ and its weights 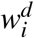 to minimise this energy allows us to derive the remaining equations in Fig. 1f. In particular, the activity level *x*^*out*^ minimising the energy must satisfy:

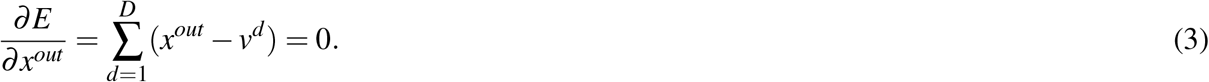

Solving the above equation for *x*^*out*^, we find the same expression as in Fig. 1f:

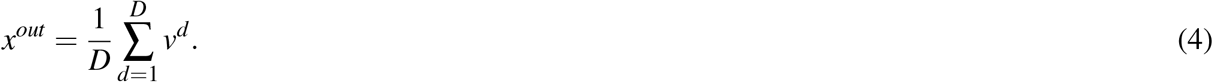

Analogously, the learning rule in Fig. 1f can be derived as a gradient descent on the energy:

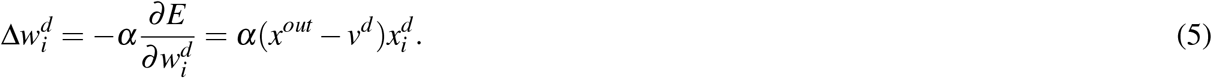

To capture properties of biological neurons and enable networks of neurons to learn effectively, we make two additional assumptions in some simulations. First, since different biological dendrites have different sizes, we assume that they may affect the somatic membrane potential *v* to different extent:

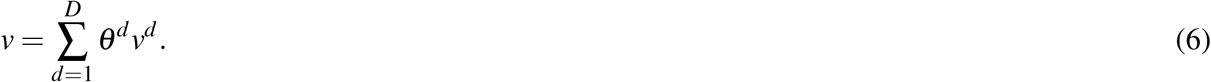

where *θ*^*d*^ denote the influence of individual dendrites, and they add up to 1. If their values are not explicitly mentioned, we use *θ*^*d*^ = 1*/D*.

Second, since the firing rate cannot be negative, we constrain the activities of neurons to be non-negative.

Thus, to capture the fact that the output from a neuron may be a non-linear function of *v*, we employ a gating notation introduced by Saxe and colleagues^63^:

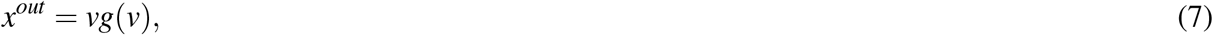

where *g*(*v*) is a gating function. If *g*(*v*) = 1, then the neuron is linear, while the rectified linear units (ReLU) can be obtained by setting *g*(*v*) = 1 when *v* > 0, and *g*(*v*) = 0 otherwise.

We show in Supplementary Information Section 4 that such neural activity (Eqs 6-7) arises from minimisation of an augmented energy function. Weight modification proportional to the negative gradient of this energy becomes:

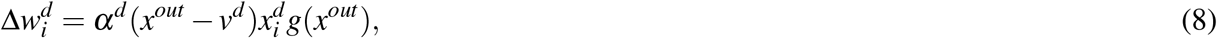

where *α*^*d*^ = *αθ*^*d*^ is a learning rate for dendrite *d*. In comparison with the weight update of the linear model (Eq 5), the above equation includes an additional term *g*(*x*^*out*^), which ensures that the synaptic weights of a neuron are only modified when this neuron is active. An analogous mechanism is also present in the backpropagation algorithm where the weight modification is gated by the derivative of the activation function^63^. It is useful to note that *g*(*x*^*out*^) = *g*(*v*) for both linear and ReLU activation functions that we employ in simulations of this paper, so the same gating is used in Eq 7 and 8. For linear neurons Eq 8 reduces to Eq 5.

### B Firing-rate formulation for networks

Neurons described by the PDM can be connected into networks, and in this paper we consider a simple case of neurons organised into *L* layers, where layers 1 and *L* are the input and the output, respectively (Fig. 1g). We denote the activity of neuron *i* in layer *l* by 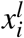. In the networks analysed in this paper, a neuron 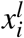 is connected to all neurons in adjacent layers, and we denote the forward and backward connections to that neuron by 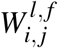 and 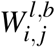, where *j* is the index of neurons in adjacent layers. We consider a simple organisation of inputs, where all forward inputs arrive at one dendrite and all backward inputs at another, and we denote the membrane potentials in these dendrites by 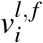 and 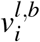, respectively.

During training the activity of input and output neurons is constrained to the training input **x**^*in*^ and target **x**^*target*^, and the activity of hidden neurons is initialised to 0. The activity of all hidden neurons is updated until it reaches equilibrium (with only 1 hidden layer, the equilibrium is reached already after the first iteration). Next, the synaptic weights are modified. This process is summarised in Algorithm 1, where column vectors are denoted with bold, and *◦* denotes element-wise multiplication.

When a network makes a prediction, only the input neurons are constrained, and the activity of other neurons is updated until convergence, as summarised in Algorithm 2.

It is important to highlight that, unlike in standard predictive coding models, there is *no global energy function* that is minimised by a PDM with respect to both the activities and weights. This is because each neuron minimises its own local energy (as defined in Eq 2), which ignores the impact of its activity on the energy of other neurons (in the same layer and in neighbouring layers). Consequently, there is no theoretical guarantee that the minimisation of neuron-level local energies leads to the minimisation of their *sum* (which can be seen as the global energy of the network). This is the fundamental reason why, in its basic form, PDMs are challenging to scale to wide and deep networks, as demonstrated in Fig. 4. While this may appear as a strong limitation of the PDM, we see it as a feature. The brain can be argued to operate under similar constraints since it is made of independent neural components, and modifying PDM to minimise the global energy would reintroduce the biological implausibilities of standard predictive coding models, such as the symmetry of forward and backward weights.

### C Firing-rate model with contrastive learning

Here we describe the “contrastive” variant of the PDM tested in Fig. 4. As in the previous section, we consider networks where each (non-input) neuron receives forward and backward dendritic predictions of its activity. For brevity, we denote the dendritic prediction errors at each layer as:

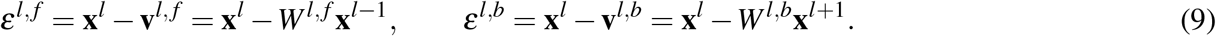

#### Algorithm 1

Learning with the PDM.

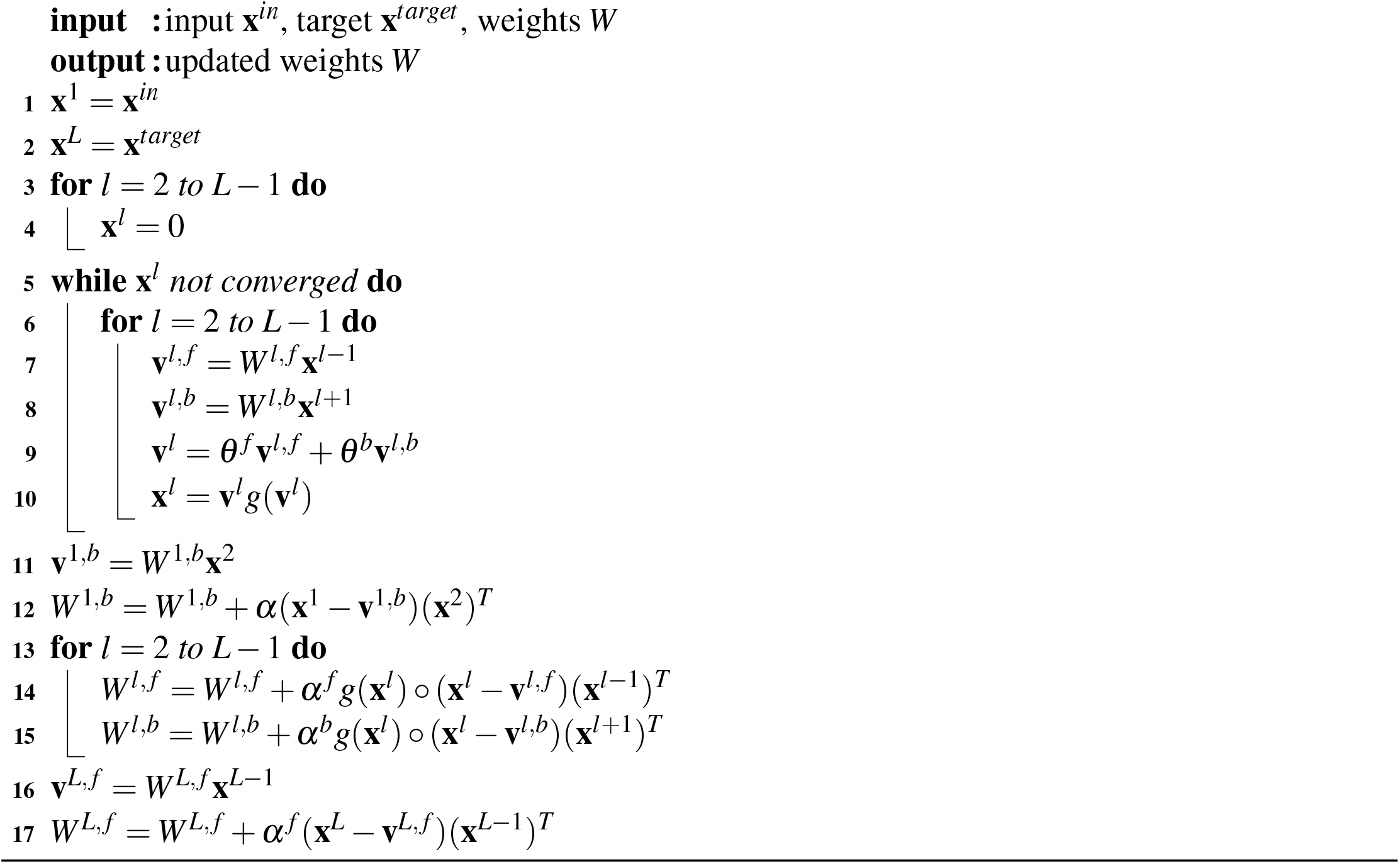

#### Algorithm 2

Prediction with the PDM.

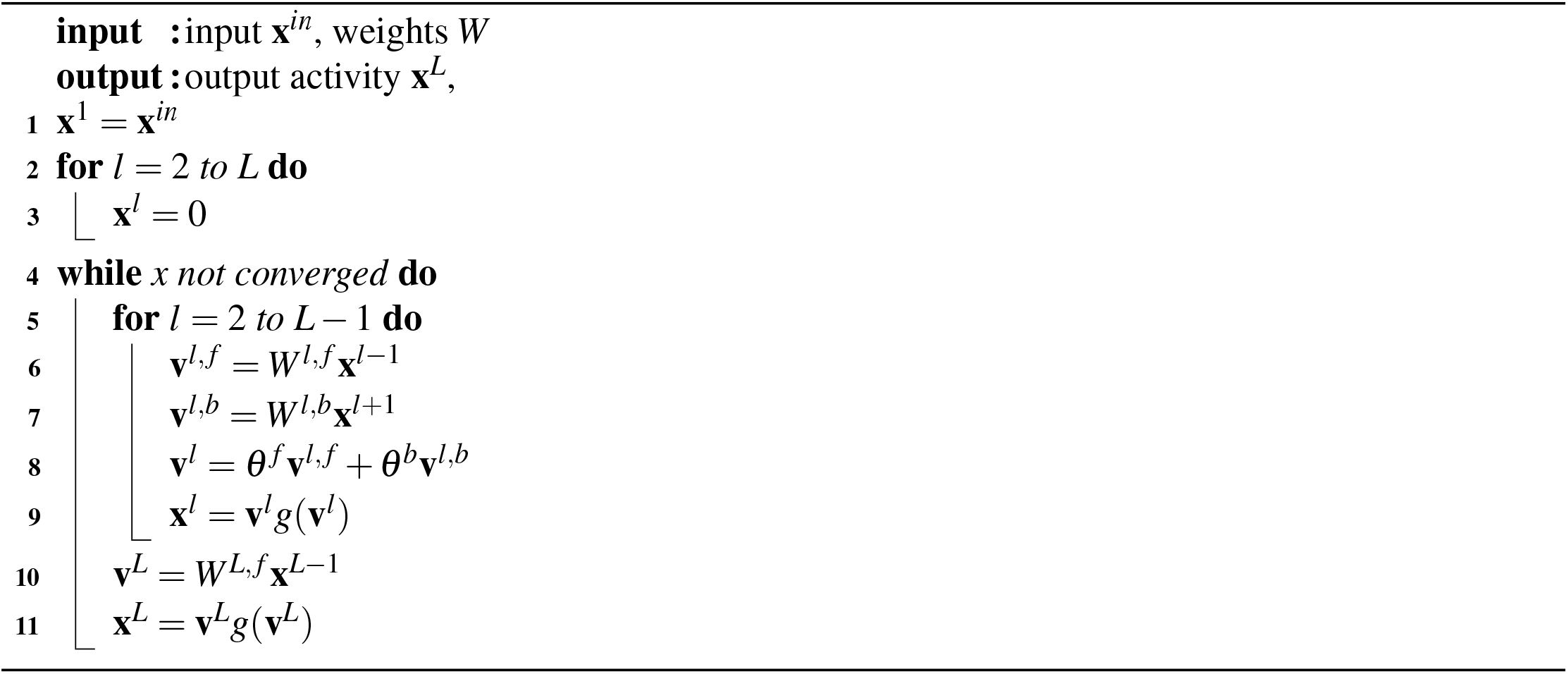

#### Layerwise energies

For each non-input layer we define a local energy. The energy of a hidden layer is given by the sum of its forward and backward prediction errors squared:

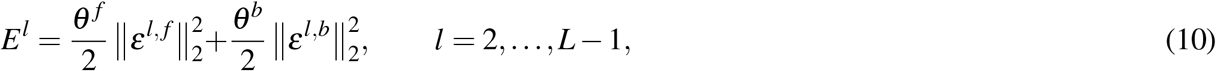

where *θ* ^*f*^, *θ*^*b*^ > 0 are fixed forward and backward gains. The output layer has no backward prediction and instead carries a supervised attraction toward the target, weighted by the nudging strength *β ≥* 0,

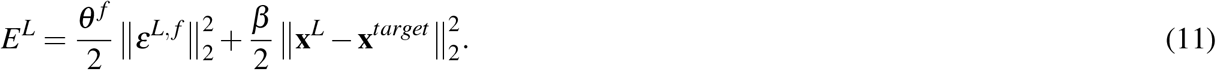

#### Neural dynamics

Activities of non-input layers are inferred by descending their local energies. At each inference step *τ*, each non-input layer is updated by gradient descent followed by an elementwise nonlinearity *f* ^*l*^,

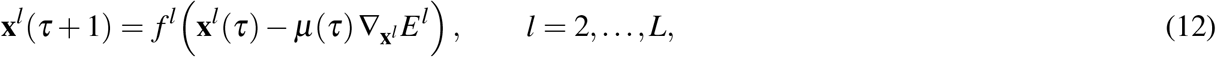

where *µ*(*τ*) > 0 is a step size that can vary during inference. The experiments of Fig. 4 use ReLU as non-linearity. Using Eqs 9-11, the gradients of each layer energy with respect to their activities are

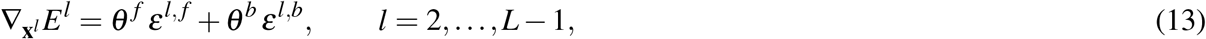

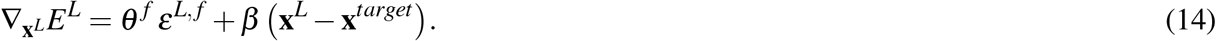

Thus, each hidden neuron is driven toward consistency with both its forward and its backward prediction, while the output neuron is additionally pulled toward the target with strength *β*.

#### Weight updates

Recall from Eq 5 that the energy gradients of forward and backward weights are given by

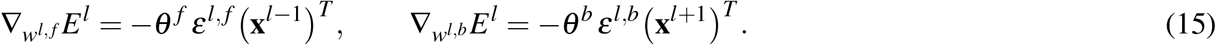

The factors (*θ* ^*f*^, *θ*^*b*^) are absorbed into the synaptic learning rates (*α*^*l, f*^, *α*^*l,b*^), respectively. The contrastive learning rule tested in Fig. 4 uses an equilibrium-propagation^48^ style update. Specifically, it contrasts these gradients between two relaxed states of the network: a free state and a nudged state. As we explain next, the two phases differ only in the output nudging strength *β*.

#### Free phase (*β* = 0)

In the free phase, the output energy (Eq 11) reduces to its forward prediction term, so all non-input layers relax under Eqs 12-14 without the supervised term. We denote the resulting free equilibrium activities and errors by 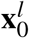 and 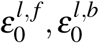.

#### Nudged phase (*β* > 0)

The network is then re-relaxed with a positive nudging factor *β* > 0, starting from the free equilibrium. Note that only the output dynamics change, receiving the supervised signal *β* (**x**^*L*^ − **x**^*target*^) in Eq 14. We denote the nudged equilibrium quantities by 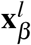 and 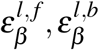.

The equilibrium-propagation-style update contrasts the weight gradients (Eq 15) between the nudged and free equilibria, giving the forward update

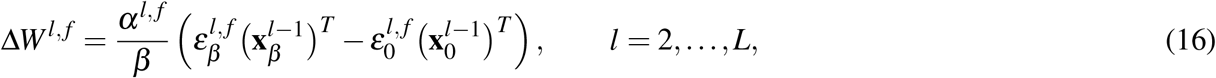

##### Algorithm 3

Contrastive learning with the PDM.

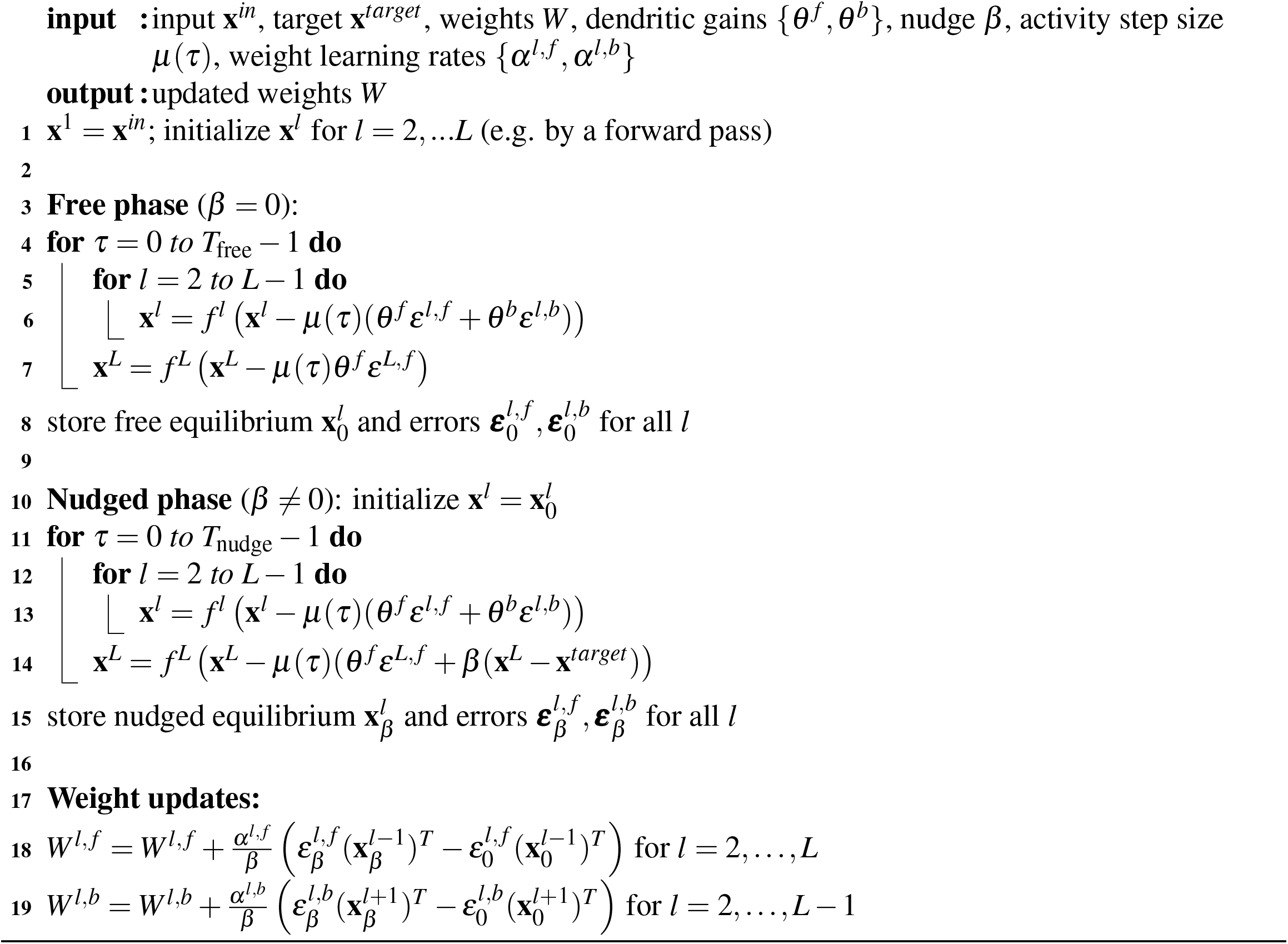

and the backward update

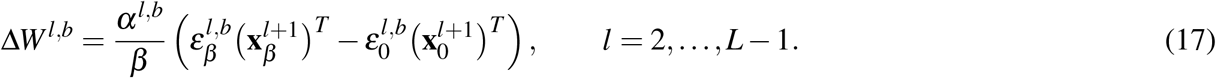

The full procedure is summarized in Algorithm 3 for a single input-output pair.

The contrastive learning rule is “local in space” as it only depends on information available to a synapse, but not “local in time”, as the information from the “free phase” needs to be stored until the target is provided. Various mechanisms have been proposed by which such contrastive update could be realized in the brain^64,65^.

### D Simulations of firing-rate models on the XOR problem

The XOR problem has two inputs, and one output target. The mapping from the two inputs 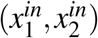 to one target is as detailed in Fig. 3a.

In Fig. 3c we trained Hebbian and predictive dendrite networks with one hidden layer and 64 hidden neurons. The Hebbian baselines had only feedforward connections, and updated weights with the simple rule 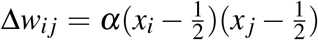 for pre- and post-synaptic inputs *x*_*i*_ and *x* _*j*_, respectively. The bounded Hebbian model clamped the weights to |*w*_*i j*_| *≤*1. For all models, the weights were initialized via standard Kaiming/He uniform initialization^66^. The networks were trained for 500 iterations, where each iteration or weight update included all the 4 possible input-output pairs for XOR. We show results for learning rate *α* = 0.05. Fig. 3c shows the mean squared error over the batch during training. Results are averaged over 8 different random seeds, with shaded bands as 95% confidence intervals.

### E Testing the firing-rate model and its extensions

All the models in Fig. 4, Supplementary Fig. 4, Supplementary Fig. 5 and Supplementary Fig. 6 were trained for 5000 steps or weight updates using a batch size of 64. Following standard practice, for each model we performed a learning rate grid search over *α* ∈ { 5 ·10^−1^, 10^−1^, 5 ·10^−2^, 10^−2^, 5 ·10^−3^, 10^−3^, 5 ·10^−4^}, selecting runs with the lowest mean validation loss (over multiple seeds) during training. For the toy linear and non-linear tasks, we tested the models on the same validation set over which the learning rate search was performed, to see if they were even capable of overfitting. For the Iris task, on the other hand, we performed a cross-validation search over 5 random folds, and report the performance on a held-out test set with an 80/20 split. Unless otherwise stated, (i) all the PDMs used an equal weighting of forward and backward errors (*θ* ^*f*^ = 0.5), (ii) both weights and activities were updated with standard gradient descent, and (iii) the standard Kaiming/He uniform initialisation^66^ was used to initialise the weights, i.e. *w*_*i j*_ *~U* (−*a, a*) where *a ∝* 1*/n* and *n* is the number of hidden units.

In Supplementary Fig. 4b, we trained one-hidden-unit linear networks (Supplementary Fig. 4a) to learn a noisy line *x*^*target*^ = 3*x*^*in*^ + *ξ*, with *x*^*in*^ *~ N* (0, 1) and *ξ ~ N* (0, 0.5). Weights were updated using the Adam optimiser^67^. In Supplementary Fig. 4f, we trained networks with two hidden units and ReLU as activation function to learn the non-linear sum 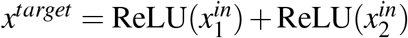, with 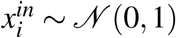. As in the XOR task, all weights (including lateral connections) were initialised using the Xavier uniform initialisation^68^. The dataset consisted of 500 examples, split into 400 and 100 for training and validation, respectively. In Supplementary Fig. 5, networks with 16 hidden units were trained to classify the Iris dataset^69^.

In Fig. 4 and Supplementary Fig. 6, we trained one-hidden-layer networks to classify MNIST and CIFAR-10 images for 15 epochs using batch size 20. The networks had 500 hidden units for MNIST and 1000 for CIFAR-10. Each dataset was split into train, validation and test sets, and all results are aggregated across 5 seeds. We tested the following models: the standard PDM, a variant of the model with contrastive learning (Section C and Fig. 4), and two backpropagation baseline models, one with frozen input-to-hidden weights and the other with fully learnable weights. For each model, we performed a learning rate sweep and selected the run with the lowest validation error, as standard practice. All models used ReLU as non-linearity. The backpropagation baselines used Adam as optimiser in Fig. 4 even though SGD was used for PDM to provide a more demanding baseline, while in Supplementary Fig. 6 performance for backpropagation with both Adam and SGD is shown for completeness. Both PDMs used error gains *θ* ^*f*^ = *θ*^*b*^ = 0.5, and the inference step size was decayed by a factor of 0.01, with a minimum step of 0.02. The standard model used 20 inference iterations. On MNIST, its weight learning rates were *α*^1, *f*^ = *α*^2, *f*^ = 0.005, and *α*^2,*b*^ = 0.001, with *µ*(*τ*) = 0.6. On CIFAR-10, they were *α*^1, *f*^ = *α*^2, *f*^ = 0.0025, and *α*^2,*b*^ = 0.0005, with *µ*(*τ*) = 1.0. The contrastive model used the same decay, with *µ*(*τ*) = 1.6 on both datasets, *T*_free_ = 30 steps in the free phase and *T*_nudged_ = 10 in nudged phase. Its weight learning rates were *α*^1, *f*^ = 0.05, *α*^2, *f*^ = 0.035, and *α*^2,*b*^ = 0.0075 on MNIST, and *α*^1, *f*^ = 0.006, *α*^2, *f*^ = 0.003, and *α*^2,*b*^ = 0.003 on CIFAR-10. For the nudged phase, we optimised for different values of *β*, showing the best results with *β* = 0.05 for MNIST and *β* = 0.0025 for CIFAR-10.

### F Spiking formulation for individual neurons

The spiking formulation of the PDM builds on several key ideas from spiking predictive coding networks^31^. In this section we first introduce the description of a single neuron, as it employs simpler notation (without multiple indices), and in a later section we give details of simulated networks.

We assume that a neuron produces spikes at discrete time points, and define the spike train *s*^*out*^(*t*) to be equal to a Dirac’s delta at time *t* when the spike is produced and 0 otherwise. Analogously we define the spikes 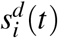 of input neuron *i* projecting to dendrite *d*. We define the signals of the neurons as:

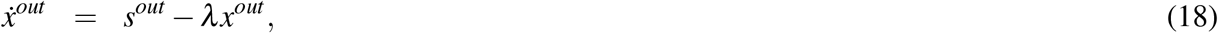

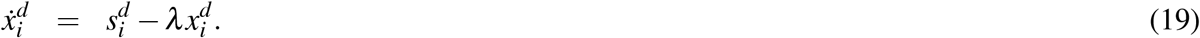

In the above equations, a dot over a variable denotes its derivative over time and *λ* is a decay parameter. Since the Dirac’s delta is defined to have an area of 1, whenever a spike is produced, the signal of the neuron increases by 1 and then decays with a time constant 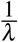. We define the membrane potential on dendrite *d* as in the firing-rate formulation (Eq. 1) but we additionally include a synaptic transmission delay *τ*:

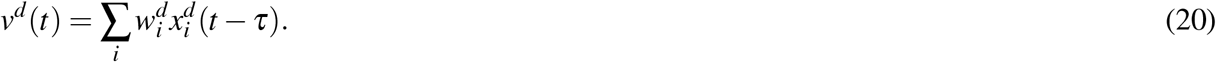

We assume that the neuron tries to minimise the same energy function as in the firing-rate formulation (Eq. 2) which we rewrite below for convenience:

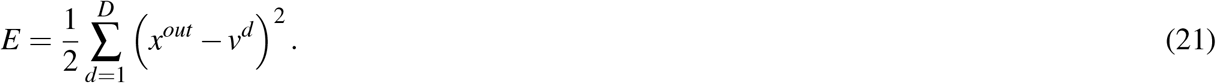

The variables in the above equation are functions of time *t*, but for brevity we omit indexing by time where we feel it does not bring additional insights. Analogously to spiking predictive coding^31^, we assume that a neuron produces a spike if and only if it reduces its energy, i.e., the spike is produced if the following inequality is satisfied:

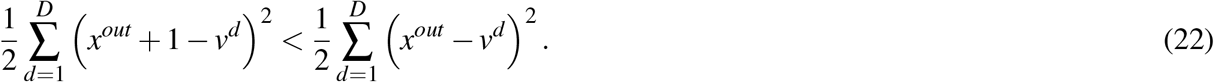

The right side of this inequality is the energy without a spike, while the left side is the energy with a spike, and it includes the term +1, because a production of a spike would increase the signal *x*^*out*^ by 1. After expanding the squares, we note that most of the terms cancel, and the inequality simplifies to:

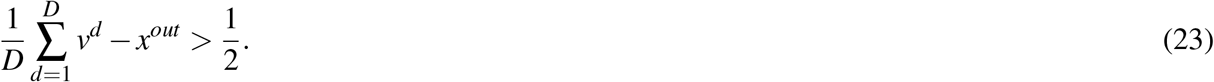

The above inequality implies that a spike should be produced whenever the left side, which depends on the average of dendritic potentials, exceeds a constant value. Hence following spiking predictive coding^31^, we interpret the left side of the inequality as the somatic membrane potential and denote it by *v*, while we interpret the right side as a spiking threshold. To see that the left side of the inequality indeed corresponds to the membrane potential, it is helpful to evaluate its derivative over time:

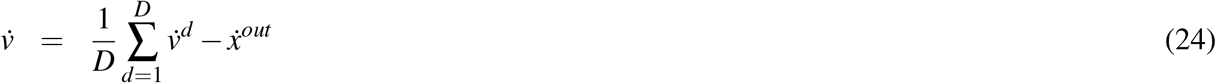

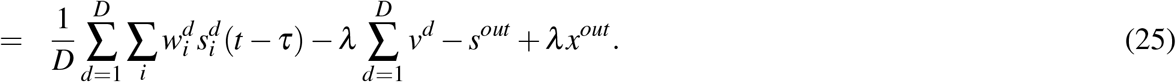

In the transition from the first to the second line, we substituted Eqs. 18-20. We observe that the second and the last terms in the second line correspond to − *λv* according to the definition of the somatic membrane potential (left side of Eq 23), hence the equation for its dynamics can be simplified to:

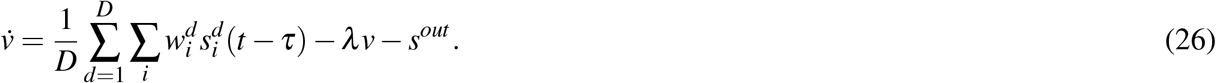

The above equation, together with the condition for generating spikes of Eq 23, correspond to the integrate-and-fire model: the membrane potential is increased by synaptic inputs, and it decays in proportion to its value, as expected from a leak current. Additionally, when the membrane potential exceeds a threshold of 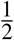, the last term in Eq 26 ensures that the membrane potential is decreased by 1, so it is reset to 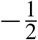. For mathematical simplicity, we keep the threshold at 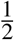 rather than at physiological values, but the model could be easily adjusted (by adding constant terms to its equations) to shift the range of membrane potentials.

The synaptic weights are modified analogously to the firing-rate formulation (Eq 8), namely, they change to reduce the energy, but they should be only be modified if the neuron is active (so that the neurons adjust representations of features they represent only if these features are present). In Supplementary Information Section 4 we define an augmented energy, and find its negative gradient over the weights, which leads to the following weight update:

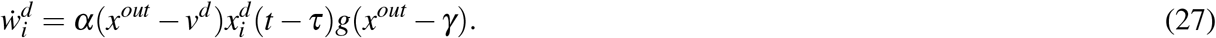

The non-linear gating needs to be modified in the spiking formulation because *x*^*out*^ remains positive after a neuron produces a single spike; hence we assume that the plasticity happens if the activity is above a small constant *γ*, namely when *x*^*out*^ − *γ* > 0. To account for imprecision in gating in biological neurons, we use a gating function equal to a steep sigmoid rather than a step function:

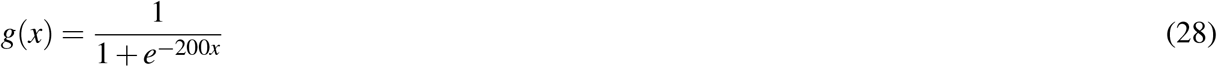

Such sigmoid gating function results in a smoother shape of an STDP curve (the STDP curve for *g*(*x*) = 1 if *x* > 0 and *g*(*x*) = 0 if *x ≤* 0 closely matches an analytic prediction shown as dashed line in Fig. 6h which has a sharp turn around −35ms). To avoid confusion, it is worth clarifying that the sigmoid shape of *g* does not imply that the firing rate of a neuron is a sigmoid function of its input (as often assumed in computational models), because here the neurons are described by the integrate-and-fire model, for which the relationship of input and firing rate is approximately a ReLU function^70^.

The spiking model of a neuron has 4 parameters, and if they are not mentioned, the following default values were used. The decay parameter was set to *λ* = 0.1, corresponding to a membrane time constant of 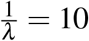 ms, which is the value was measured at body temperature from layer 5 neocortical neurons^71^. The synaptic transmission delay was set to *τ* = 5 ms, since it is within a range of observed delays between the activity of a presynaptic neuron and the peak of a postsynaptic potential^72^. The default value for the learning rate is *α* = 0.01, while the threshold postsynaptic activity for plasticity is *γ* = 0.02.

### G Analytic formula for STDP curve

In order to compute the total weight change during a simulation of an STDP experiment, we need to compute an integral of instantaneous weight change (Eq 27) over the simulated period. We restate the expression for instantaneous weight change for convenience, and for simplicity we omit indexing of input neurons (as only one of them is active):

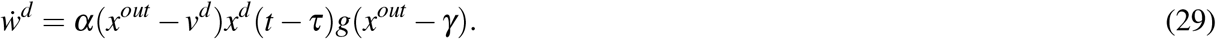

An integral of Eq 29 can be derived under assumption that the gating function *g* is a step function (rather than sigmoid):

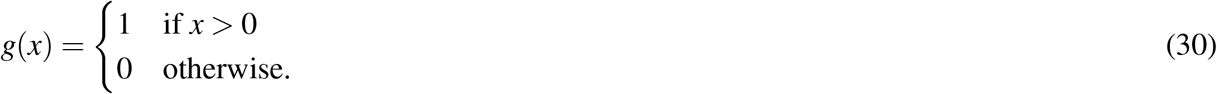

According to Eq 29, weights are modified when the postsynaptic signal is sufficiently large (*x*^*out*^ > *γ*) and presynaptic spike-impact is also non-zero (*x*^*d*^ > 0). Hence let us establish the start and end of the period when these conditions are satisfied, which will serve as limits of our integration.

Without the loss of generality, let us assume that the postsynaptic neuron produces a spike at time 0, hence:

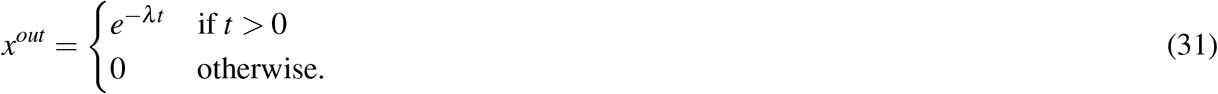

Let us denote the interval between pre- and postsynaptic spikes by *I*_*pp*_, hence the presynaptic neuron produces spikes at time −*I*_*pp*_, and its delayed input signal (occurring in Eq 29) is:

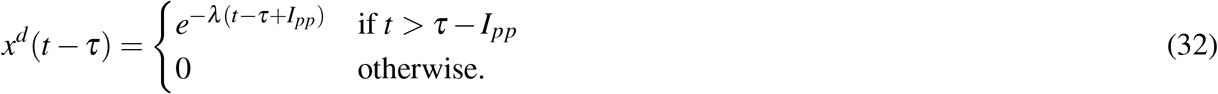

Both pre- and post-synaptic signals will have non-zero values from *T*_*s*_ = max(0, *τ* −*I*_*pp*_), hence we will start integration at this time. The gating function *g*(*x*^*out*^ −*γ*) of Eq 30 will become zero when *e*^−*λt*^ = *γ*. Hence the plasticity will end at 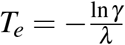.

To calculate the total weight change over the simulation, we integrate the instantaneous weight change of Eq 29:

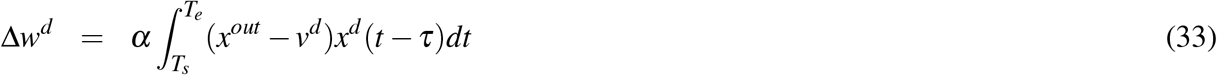

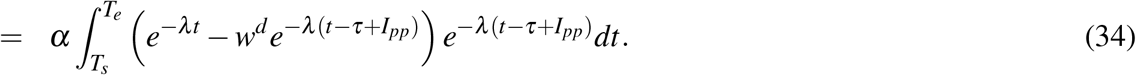

The initial weight *w*^*d*^, which appears in the second line, evolves during the simulation, but since we assumed a limit of small learning rate, it does not change sufficiently, so we ignore terms connected with its change (which are of order (*O*(*α*^2^)). Multiplying out the terms in the bracket, we obtain:

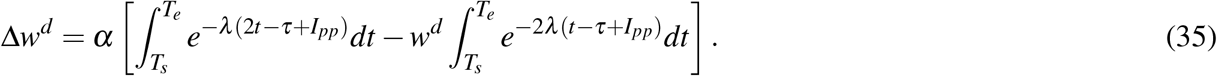

To evaluate the first integral we substitute *s* = 2*t* − *τ* + *I*_*pp*_. Differentiating this definition we get 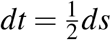. To evaluate the second integral we substitute *q* = 2(*t* −*τ* + *I*_*pp*_), and 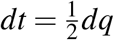. After substitution we obtain:

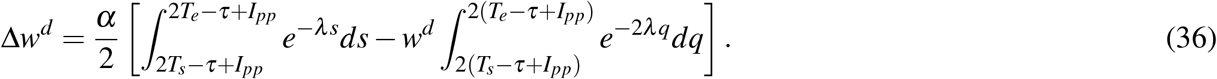

Evaluating the integrals, we obtain the final approximate expression for the STDP curve:

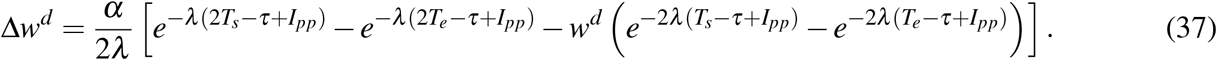

From Eq 37 one can infer the effects of parameters on the shape of STDP curve seen in Fig. 6d-g. The learning rate *α* only scales the entire equation, so it stretches the curve vertically. The decay rate *λ* scales the entire equation and scales all terms *I*_*pp*_, so it stretches the curve both vertically and horizontally. The synaptic delay *τ* is always subtracted from *I*_*pp*_, so it shifts the curve horizontally. Finally, the gating threshold *γ* affects *T*_*e*_ so it has more complex effects.

### H Fitting to STDP profile data

Fig. 6i and Fig. 6j show STDP curves from the PDM fitted to data from Figure 3 in^33^ and Figure 2 in^34^, respectively. To present the data from^33^ as a percentage of weight change in Fig. 6i, the original values from^33^ were reduced by 1 and then multiplied by 100. Values in^34^ are already presented as the same unit in Fig. 6j, so no transformation was needed. The fitting optimized the following parameters of our model in the following ranges: *τ∈* uniform(0.0, 10.0), *λ ∈* loguniform(0.005, 0.5), *γ ∈* uniform(−0.5, 0.5), *α∈* loguniform(0.01, 1.0), initial *w∈* uniform(0.0, 2.0). Fitting was performed with OptunaSearch https://optuna.org and sampled for 1000 times. The optimization objective was to minimize the squared difference between experimentally observed and predicted weight change, summed across all data points. The variability of experimental data points depends on the interval between pre and postsynaptic spikes. For example in Fig. 6i, this variability is high around 0 delay and low for larger delays. To take this variability into account, each data point (Δ*t*_*i*_, Δ*w*_*i*_) was weighted within the sum in the objective function by the reciprocal of the variance of the values Δ*w*_*j*_ among data points with delays within a window Δ*t* _*j*_ *∈* (Δ*t*_*i*_ − 5*ms*, Δ*t*_*i*_ + 5*ms*).

In Fig. 7b the parameters of the spiking PDM were fit to data describing the effect of initial connection strength on plasticity, obtained from Figure 5 in^3^. Fitting was done with OptunaSearch https://optuna.org, sampled for 1000 times, with the objective function being the sum of squared error between the weight change in the model and in the experiment data. The fitting tuned the parameters of our model in the following ranges: *τ ∈* uniform(0.0, 10.0), *λ ∈* loguniform(0.005, 0.5), *γ ∈* uniform(−0.5, 0.5), *α ∈* loguniform(0.01, 1.0). We also tuned two additional parameters that describe a linear mapping from the units of initial connection strength in the data (the initial EPSP amplitude measured in pA) to that in our model. The first parameter *w*_starting_ describes the synaptic weight in the units of our model corresponding to the starting value of the x-axis in Fig. 7b, which we denote 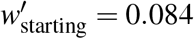 pA. The second parameter *w*_range_ describes the range of synaptic weights on the x-axis in Fig. 7b in the units of our model, corresponding to the range on the plot of 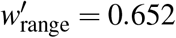 pA. These two additional parameters were tuned in the following ranges: *w*_starting_ *∈* uniform(0.0, 0.3), *w*_range_ *∈* uniform(0.0, 0.3). For given values of *w*_starting_ and *w*_range_ considered by an optimizer, the parameters of linear mapping *k* (slope) and *b* (intercept) between pA units of the model were found by solving analytically the set of equations: 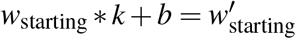 and 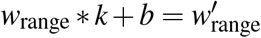.

To evaluate the ability of the spiking PDM to replicate plasticity outcomes under spike triplet paradigms, model parameters were optimized to fit the experimental data reported in^39^. The fitting minimized the sum of squared errors between model predictions and experimentally observed mean weight changes across eight triplet and single-pair conditions. Optimization was performed using `fmincon` in MATLAB (MathWorks) with `MaxFunctionEvaluations` = 10000 and `TolFun` = 10^−8^. To avoid local minima, 50 random initializations were used and the solution with the lowest SSE was selected. The induction protocol was simulated as 60 repeated cycles at 1 Hz, consistent with the experimental procedure. The following parameter bounds were imposed: *τ ∈* uniform(0.0, 10.0), *λ ∈* loguniform(0.01, 0.5), *γ ∈* uniform(−0.5, 0.5), *α ∈* uniform(0.001, 2.0), *w*_starting_ *∈* uniform(0.0, 1.0).

### I Spiking formulation for networks

Spiking neurons described in the previous section can be connected into networks analogously as for firing-rate formulation, and in this paper we consider a simple case of neurons organised into layers, and receiving forward and backward inputs on separate dendrites (Fig. 1g). In spiking models of such networks we index all variables introduced in the previous section analogously to how we did this in the firing-rate formulation. The training input **s**^*in*^ and target **s**^*target*^ are now both spike trains. Each training iteration has duration *T*, and in our simulations the time is divided into discrete intervals Δ*t*. Thus the training patterns **s**^*in*^ and target **s**^*target*^ are matrices of dimensions: number of neurons in the corresponding layer by 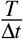. They contain 1 at positions corresponding to times when an input or target neuron produces spikes, and 0 elsewhere. Since the spiking model includes transmission delays, the membrane potentials are being integrated from time *τ* so that their values are defined. The simulation procedure during a single training iteration is summarised in Algorithm 4, where for brevity the comments specify for which indices particular operations are applied. The simulation procedure during prediction is summarised in Algorithm 5.

### J Simulations of the spiking model in the XOR problem

Training on a single pattern of inputs and target consisted of a simulation of *T* = 50 ms of neural activity. Each training pattern (**x**^*in*^, *x*^*target*^) from the XOR problem (Fig. 3a) was encoded as sequence of spikes during a single simulation. If a particular input or target was 0 for a given pattern in the XOR problem, the corresponding training sequence did not contain any spikes, *s*_*i*_(*t*) = 0 for all *t*. If, on the other hand, an input or target was 1 in XOR, the corresponding *s*_*i*_(*t*) contained a regular sequence of 5 spikes with intervals equal to 10 ms. These intervals were chosen such that they were equal to twice the value of the transmission delay *τ* = 5 ms, so that spikes on an input may result (after a transmission through 2 layers)

#### Algorithm 4

Learning with the spiking PDM.

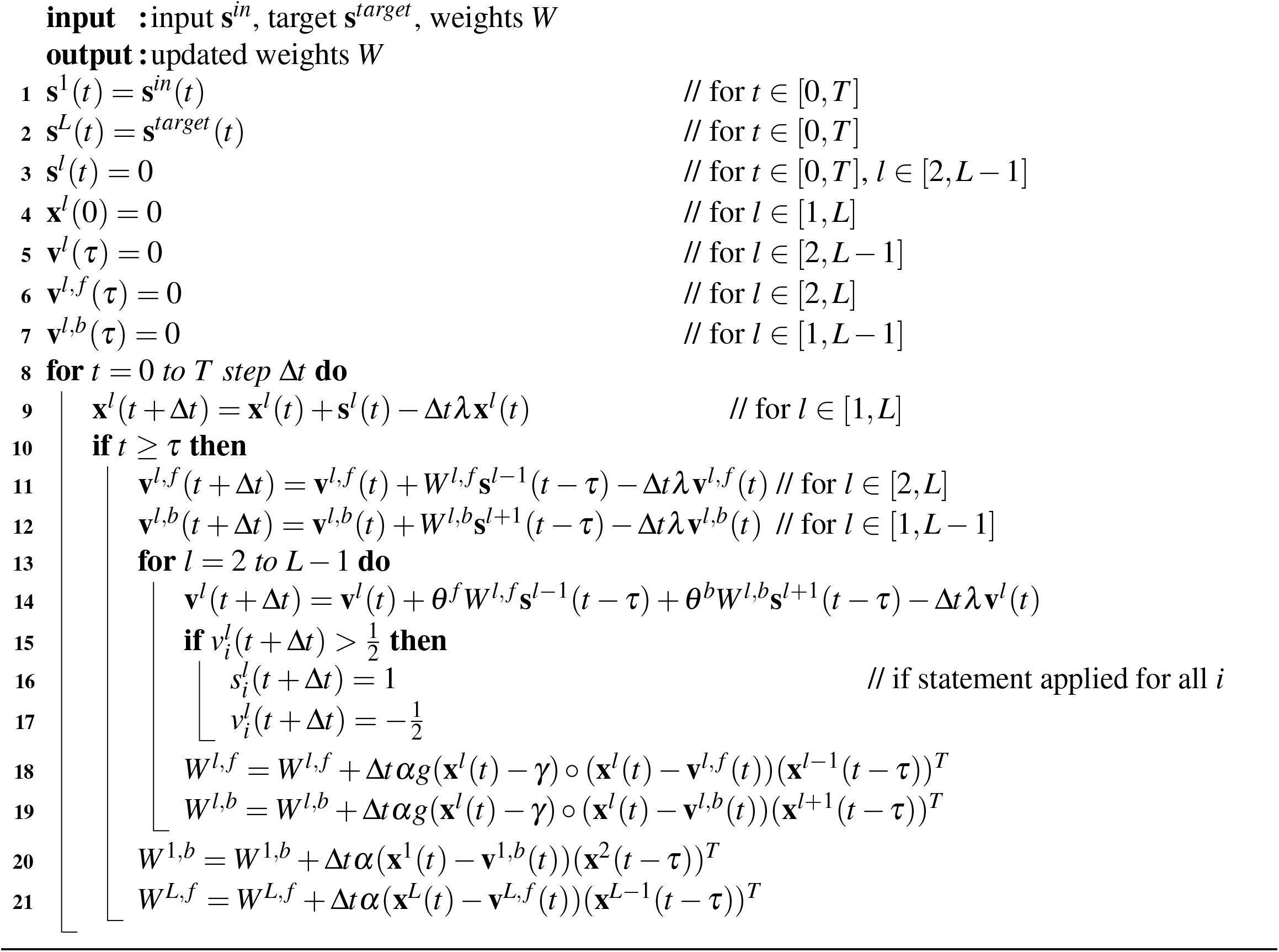

#### Algorithm 5

Prediction with the spiking PDM.

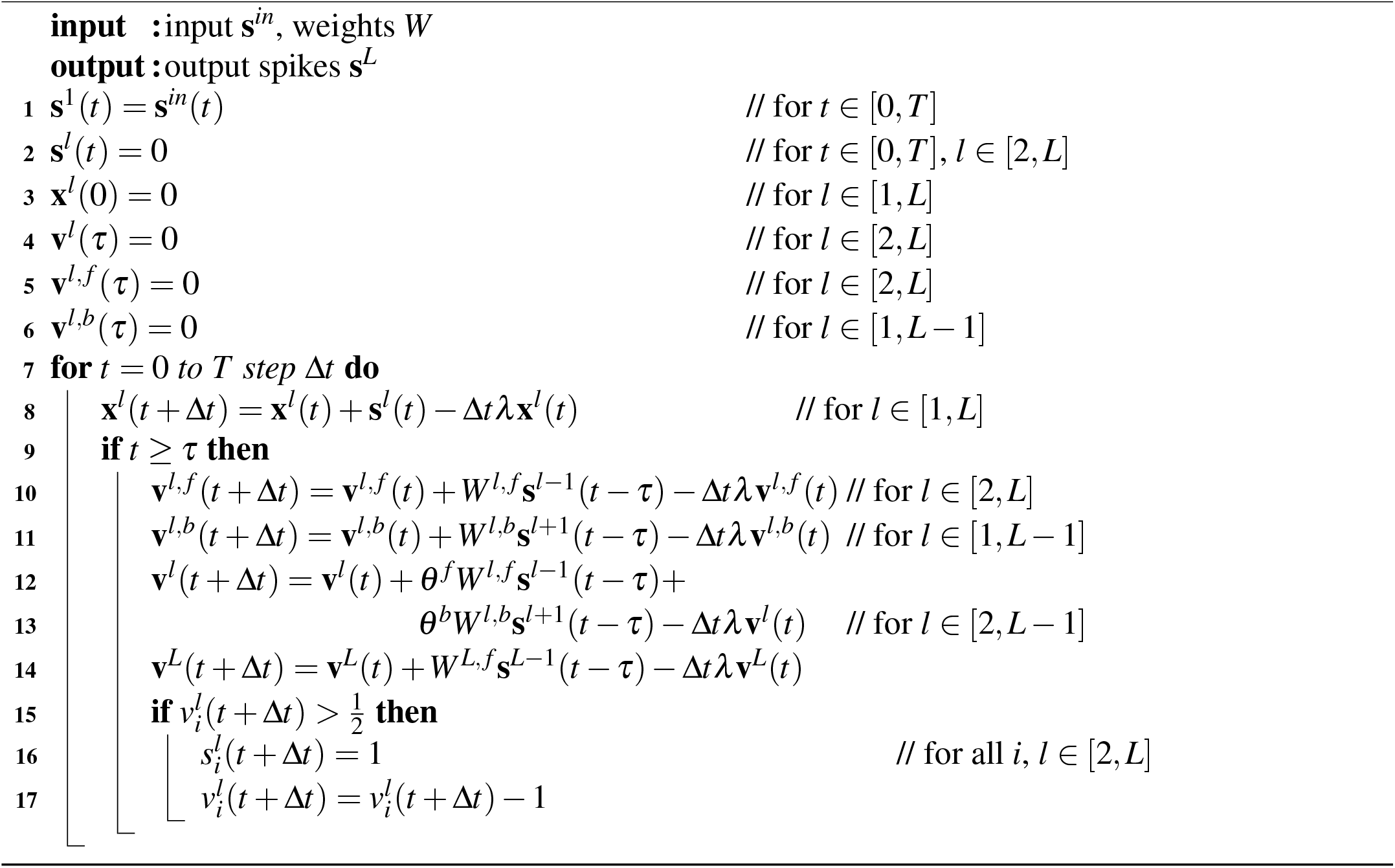

in a spike on the output coinciding with the target spike. The first spike in the sequence occurred at time *t* = 0.

During testing, only the input neurons are fixed, while hidden and output neurons are driven by their synaptic input. The network could produce up to 4 spikes, due to the transmission delays, hence the training error for a single example was defined as 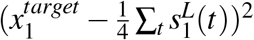. Fig. 3d plots the mean squared error over all possible (4) examples during training. Results are averaged over 10 different random seeds, with shaded bands representing one standard deviation.

We simulated a spiking PDM and a classic online STDP model^22^ with one hidden layer containing *N*_2_ = 128 neurons. For all models, the forward weights to the hidden layer *w*^2, *f*^ were initialised from a uniform distribution with limits − 2 and 2, the forward weights to the output neurons were initialised to 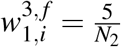, while the backward weights to the hidden neurons were initialised to 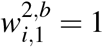. Across all experiments, we used an integration step Δ*t* = 0.1 ms and learning rate *α* = 0.005. For the PDM, the parameters describing the influence of forward and backward projections on the activity of hidden neurons were set to *θ* ^*f*^ = 0.6 and *θ*^*b*^ = 0.4, to ensure that the forward input can produce a spike during prediction.

For the classic STDP model^22^, neurons are implemented as leaky integrate-and-fire units: synaptic input is integrated into a leaking membrane potential *v*_*i*_ (membrane time constant *τ*_*m*_ = 10 ms), and a spike is generated when *v*_*i*_ reaches a threshold of 0.5, followed by reset to −0.5. Each presynaptic neuron maintains a trace 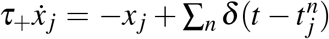, and each postsynaptic neuron maintains a trace 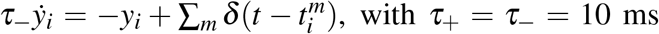. The weights are updated with the simple rule 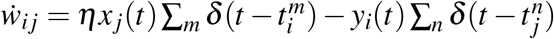 with learning rate *η* = 0.005, giving potentiation when the postsynaptic neuron fires shortly after presynaptic activity, and depression when a presynaptic spike arrives while a postsynaptic trace is elevated.

### K Statistics and Reproducibility

The work in this paper involved computer simulations, but due to random initialisation of weight parameters, the simulations were repeated multiple times. No statistical method was used to predetermine the number of repetitions. No data were excluded from the analyses. Since the order of execution has no effect on the result of numeric experiments, they were not randomised. The investigators were not blinded to outcome assessment.

To visualise the variability of simulation results, we either presented individual data points, or error bars showing confidence intervals. Confidence intervals were computed using bootstrap throughout the paper and detailed description of the implementation can be found in https://seaborn.pydata.org/tutorial/error_bars.html#confidence-interval-error-bars.

### L Use of Large Language Models

ChatGPT was asked for feedback on an earlier version of the manuscript, and we made several adjustments to the manuscript’s organization and wording based on the obtained feedback.

## Data availability

Learning tasks were built using the publicly available Iris^69^, MNIST^73^ and CIFAR-10^74^ datasets. They are available at https://archive.ics.uci.edu/dataset/53/iris, https://web.archive.org/web/20200430193701/http://yann.lecun.com/exdb/mnist/ and https://www.cs.toronto.edu/~kriz/cifar.html, respectively.

## Code availability

Complete code and full documentation reproducing all simulation results written in Python is made publicly available at https://github.com/Bogacz-Group/PDM. It is released under the MIT license.

## Supplementary Figures and Information

We describe here analyses of a linear version of the PDM, which provide theoretic grounding for its ability to perform credit assignment. We start with presenting the properties of synaptic weights, activity and prediction error in the model, which we will use in subsequent analyses. Then, we demonstrate that the PDM modifies weights in a whole network, such that its output moves directly towards the target. Afterwards, we identify conditions under which the model has the same dynamics as a variant of predictive coding. Next, we define and analyse an energy function for a non-linear version of the model. At the end, we analyse the reasons for the representation collapse in the PDM.

**Supplementary Fig. 1.**
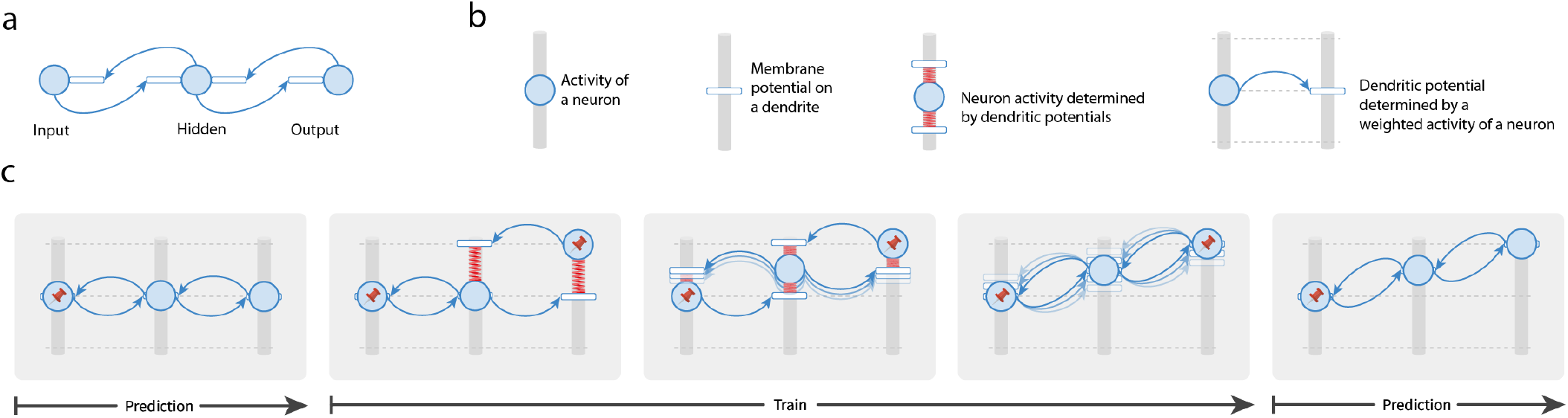
The mechanical analogy of the PDM illustrating that when an error occurs at the output layer, the network can propagate the error through the network. ▶ **a** |Example of a simple network with 1 neuron per layer, for which a mechanical analogy is presented. ▶ **b**| Components of an abstract machine with the same dynamics as the network in panel a. In the machine analogy the activity of each neuron corresponds to the height of a node (represented by a blue circle) sliding on a vertical post. The membrane potential on each dendrite also corresponds to the height of a sliding node (represented by a wide rectangle). The node corresponding to a neuron’s activity can be thought of as connected to the nodes corresponding to dendritic potentials by springs that try to bring them together (indicated in red). Since the membrane potential depends on the activity of neurons projecting to this dendrite, the arrows denote the transfer of information. The extension of a spring corresponds to a dendritic error, and the potential energy of all the springs associated with a neuron is proportional to the sum of squared dendritic errors, which we refer to as the energy of a neuron. ▶ **c** | Changes in the configuration of the machine during learning. To understand how the PDM propagates errors during learning, consider the changes in the configuration of the machine during training on a single example. Assume that before training all the weights are equal to 1 (so that the dendritic potential is equal to the activity of a neuron projecting to this dendrite; first display). When the left neuron is fixed to an input (indicated by a pin in the first display), adjusting the activity of the other neurons removes any errors, and the network produces an output equal to the input. Providing a target output pattern corresponds to changing and fixing the activity of the output node (represented by a pin in the right node in the second display). Importantly, the change in the output activity will also result in a change in the activity of the hidden neuron (as it is pulled up by a spring from one of its dendritic nodes in the third display). Furthermore, due to this change in configuration of neural activity, the springs are extended across all layers (third display), implying that the error on the output has been propagated throughout the network. Once the activity stabilises, the weights throughout the network are modified to further reduce the energy (fourth display). With such modified weights, the network can generate an output consistent with the training input (fifth display).

**Supplementary Fig. 2.**
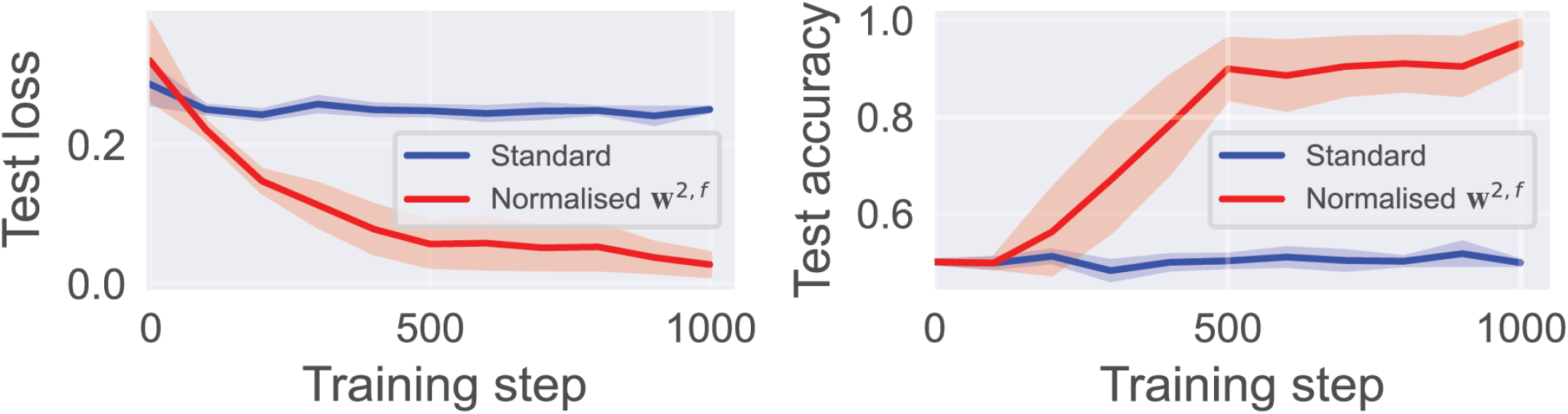
Learning XOR with a different non-linearity in the PDM. We investigated a version of the PDM where the non-linearity is applied on each dendrite, so that each dendritic potential becomes 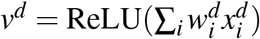. During training on a particular example, the activities of hidden neurons are found which minimize their energies (the activity of each hidden neuron is updated proportionally to the negative gradient of its own energy, Eq 2). Once the activity converges, the weights are modified proportionally to a negative gradient of the energy, which now becomes: 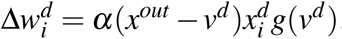. This weight update looks similar to that used in the main text (Fig. 3b), but it is gated by *g*(*v*^d^) rather than *g*(*x*^*out*^), and gating by *g*(*x*^*out*^) is critical for producing STDP. ReLU networks with 2 hidden units were trained on the XOR task for 1000 steps, using Adam, batch size 64 and learning rate *α* = 0.005. During both training and testing on each example, the activities of hidden neurons were found by updating them with gradient descent (proportionally to the negative gradients of neurons’ energies) for 20 iterations with step size 0.5. Plotted are the test loss (*left*) and accuracy (*right*) over training for models with and without normalised forward weights. Solid lines and error bars indicate means and standard errors over 10 random seeds.

**Supplementary Fig. 3.**
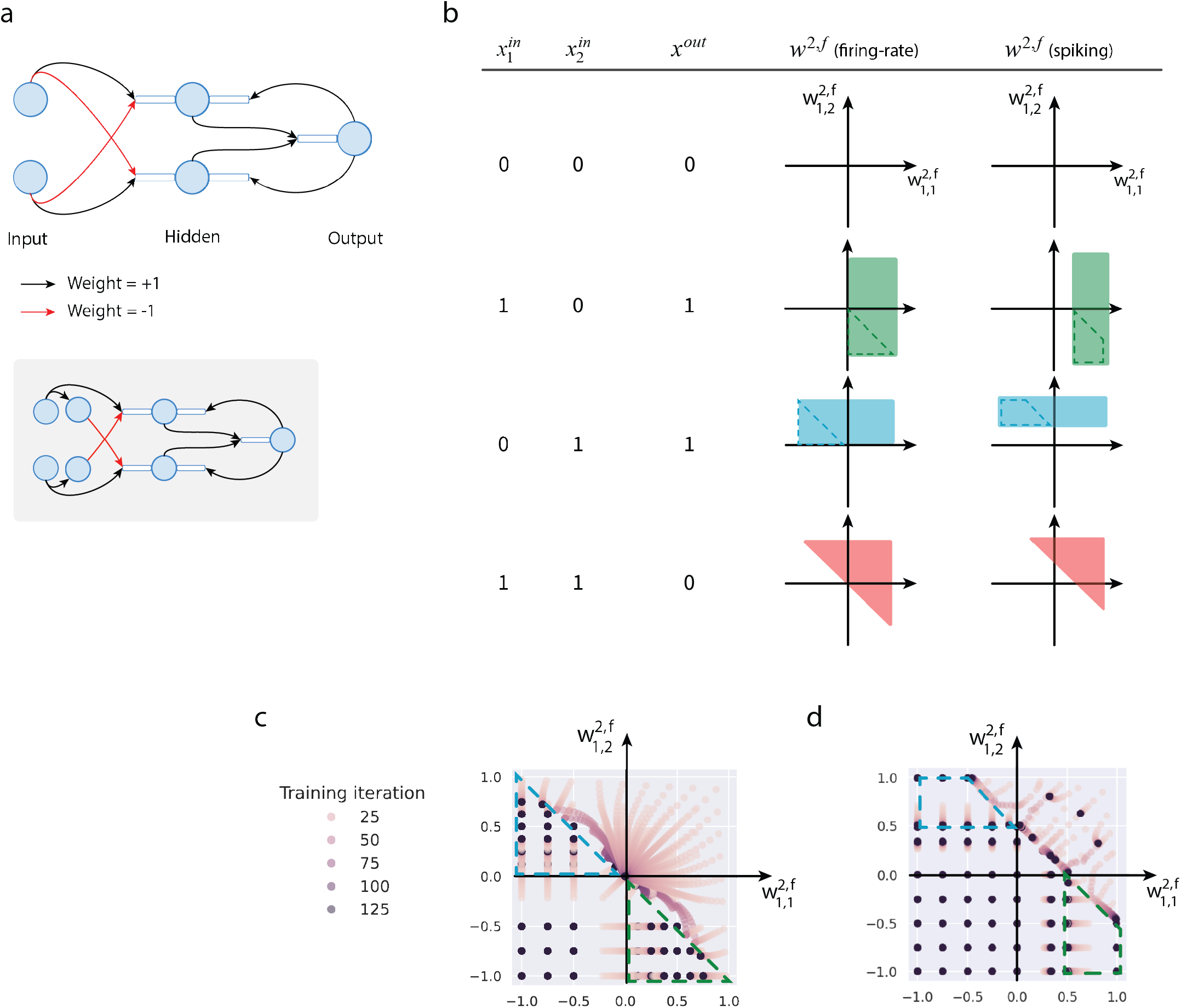
Representation learning in the XOR problem. ▶ **a**| Configuration of a minimal PDM for solving the XOR problem. Inset shows an analogous network satisfying Dale’s law. Considering this network helps to understand what features are necessary to solve the XOR. Its hidden neurons represent two features corresponding to the training examples (1,0) and (0,1). In particular, the top hidden neuron is active only for (1,0), but not any other examples, and the bottom hidden neuron is active only for (0,1). When such neurons are present, the output neuron can sum their activity and produce the correct output. ▶ **b**| Required weight combinations of hidden neurons. To get an insight into the learning of hidden neurons, the table shows the regions of the space of forward weights required to activate a hidden neuron for corresponding training examples. Specifically, the fourth column show weights where a hidden neuron has a positive forward potential in the firing-rate model, and the last column shows the weights for which the neuron produces spikes in the spiking model (these regions are similar to those in the firing-rate model but shifted because the weights from active inputs must add up to at least 0.5, so that the membrane potential reaches the spiking threshold of 0.5.). To represent the features described in panel b, the neurons should have their weights either in the region exclusively activated by (1,0) (green dashed region), or in the region only activated by (0,1) (blue dashed region). ▶ **c**| Changes in the forward weights of a hidden neuron during training of the firing-rate model. To investigate whether the weights converge to the regions representing useful features, this panel shows the changes of the forward weights of a hidden neuron depending on its initial weights (similar to analyses of backpropagation^63^). Dots show the values of weights over training with iteration indicated by colour. In these simulations, the output-to-hidden weight is initialized to 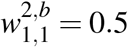. Hence, when both forward weights are initially below −0.5, this neuron is never activated during training so its weights do not change; such neurons help the network to produce the output 0 for input (0,0). When all the initial forward weights are positive, the neuron is most strongly activated by the last training example from Fig. 3a, but the target output for this example is 0, so the weights converge to 0; such neurons help the network to produce output 0 for input (1,1). Importantly, there exist large regions in the space of initial weights where the forward weights converge to values encoding useful features for computations of XOR, represented by blue and green dashed triangles in panel b, and the hidden neurons with such weights enable the networks to produce output 1 for inputs (1,0) and (0,1). The methods used for these simulations were as follows. We train a network with one hidden layer, and one hidden neuron only. We set the initial value of the weight from input 1 and 2 to the one hidden neuron to different values corresponding to co-ordinates in the panels, and observe the evolution of the two weights over iterations. Learning rate is set to *α* = 0.1. Weights are updated for 128 iterations over the whole dataset. Consequently, since the dataset has 4 data points, the weights are updated for 512 iterations. The parameters describing the influence of forward and backward projections on the activity of hidden neurons were set to *θ* ^*f*^ = 0.5 and *θ*^*b*^ = 0.5. ▶ **d**| Changes in forward weights of a hidden neuron during training of the spiking model. The spiking network modifies the weights between input and hidden neurons analogously to the firing-rate model. Namely, when both weights are initially low, they do not change; if both are initially high, they decay; and importantly, there exist ranges of initial weights, where the weights converge to the desired regions where the neurons are activated by an input that should produce the output of 1.

**Supplementary Fig. 4.**
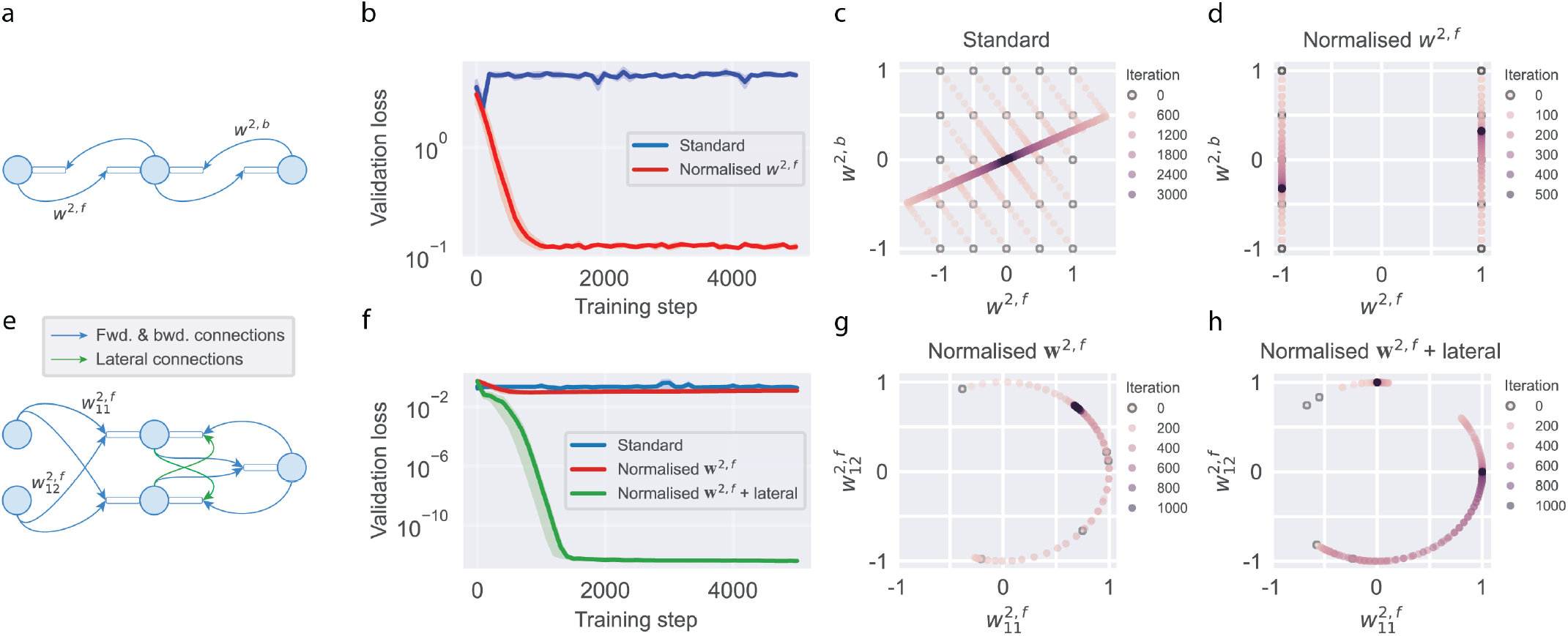
Preventing trivial weight configurations in the PDM. ▶ **a**| The top row of panels illustrates a problem occurring in the basic version of the PDM, namely that the weights feeding into a hidden neuron can converge to 0, since such weights guarantee no dendritic error and hence minimize their local energy. However, with zero weights the hidden neurons do not transmit any information, and the network cannot solve the task. We found that such collapse happens even in a surprisingly simple linear network with 1 hidden unit learning a 1-dimensional linear mapping between input and output when the training examples are noisy (*x*^*target*^ = 3*x*^*in*^ + *noise*). ▶ **b**| Validation loss of the PDMs trained on the noisy linear task. Solid lines and shaded regions indicate the mean and standard error over 5 random seeds. Although the loss of the standard model initially decreases, it then increases, because the input-to-hidden weights converge to 0 (illustrated in panel c). One way to address this problem is to normalize the weights, so that the sum of squared weights on a forward dendrite is equal to 1 (in the network with one hidden neuron, this corresponds to setting the weight to its forward dendrite to 1). With such normalization, the weights no longer converged to 0 (illustrated in panel d), and the network learned the task. ▶ **c**| Learning dynamics of the forward and backward weights (*w*^2, *f*^, *w*^2,*b*^) for the standard model trained on the toy linear task, showing collapse to 0. Coloured dots indicate weight values at a specific step during training, with the first step highlighted by a black border. Supplementary Information Section 5 provides a mathematical analysis revealing why the weights converge to 0. ▶ **d**| Same as the previous panel (c) for the model with normalised forward weights, showing no collapse to 0. ▶ **e**| The bottom row of panels illustrate a tendency to converge to another trivial configuration in which all the hidden neurons encode the same information. This happens because each hidden neuron minimizes its own local energy (which is not affected by the activity of other neurons within the layer) and hence may find the same minimum solution as other neurons. To illustrate this issue, we trained non-linear models with two hidden neurons on a task where the desired output is a sum of non-linearly transformed inputs 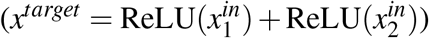. ▶ **f** | Validation loss of three different versions of the model trained on the non-linear sum task. The models that included only blue connections in panel e did not fully solve the task (the model with normalised forward weights slightly outperformed the standard model). The weights of different hidden units collapse to the same representation (illustrated in panel g). In an attempt to address this other form of collapse, we added learnable lateral connections between hidden units (green arrows in panel e). With the lateral connections, the hidden neurons learned features required for the task (illustrated in panel h) and the model learned the non-linear task.▶ **g** | Learning dynamics of the forward weights feeding into one hidden unit 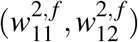 for the model with normalised weights trained on the toy non-linear task. The weights collapse to a single weight configuration encoding the sum of inputs because such activity best matches the potential in the backward dendrite receiving the output activity. ▶ **h**| Same as the previous panel (g) for the model with both normalised forward weights and lateral connections, showing weights converging to two configurations enabling hidden neurons to encode ReLU 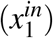 and ReLU 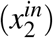. To understand why the lateral connections enable learning of the task, it is helpful to consider the configuration of weights in a network that allows the task to be solved. The forward weights of the hidden neurons need to enable the hidden neurons to encode ReLU 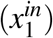 and ReLU 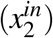, as in the panel, e.g. 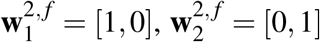, while the forward weights of the output neuron need to enable summation, e.g. **w** ^3, *f*^ = [1,1]. However, even with such forward weights there will be always prediction error during training if the lateral connections are not included. This happens because the output has less information than the input (output is 1-dimensional while input is 2-dimensional), hence the feedback can never match richer forward input. When both green connections in panel e have weights equal to − 1, there is no prediction error during training for the forward weights mentioned earlier, so such weights are the minimum of energy, and the neurons can converge to them. Intuitively, the lateral connections enable backward dendrites to have richer information - not just based on the output but also on other hidden neurons - so the backward dendrites can match forward dendrites. One could also view the role of these lateral connections as enabling the neurons to match their representation of sensory input with the target and the representations independently learned by other neurons, which is similar to a mechanism known to reduce representation collapse^75^.

**Supplementary Fig. 5.**
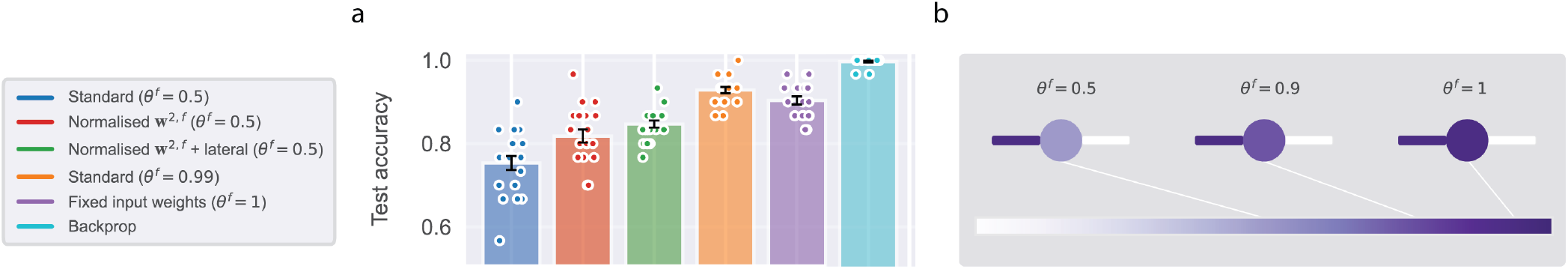
Learning the Iris task with the PDM. To probe the limits of the PDM with normalised weights and lateral connections (Supplementary Fig. 4), we benchmarked its performance in classifying species of iris^69^, a standard machine learning task. As additional controls, we tested backpropagation and a model with fixed input-to-hidden weights. We found that the model with lateral connections still provides significant improvements over the simpler model versions, but it lags behind the fixed input weights model (panel a). Encouragingly, we observed that a version of the standard model where the hidden activity is weighted much more strongly by the forward (vs backward) dendrite (*θ* ^*f*^ = 0.99, see panel b and Methods Section A) outperforms the fixed input weights model (panel a), implying that it can learn useful hidden representations. Such differential weighting of inputs could occur from the backward dendrite being more distal than the forward dendrite^76^. The stronger influence of forward dendritic predictions results in hidden neurons encoding more information about the inputs, and provides another simple mechanism preventing representation collapse. Nevertheless, none of the variants of the PDM tested in this figure match the performance of backpropagation. In panel (a), we plot the average test accuracy for one-hidden-layer networks. Error bars indicate the standard error over 20 random weight initialisations (superimposed as dots).

**Supplementary Fig. 6.**
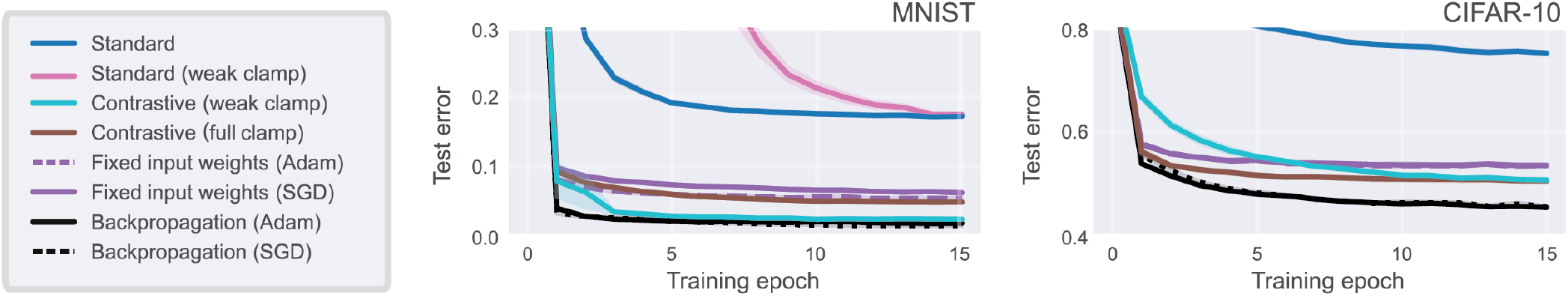
Performance of additional models and ablated versions of the models on the benchmark tasks. Standard models use a single inference phase (no free phase). The “standard” model fully clamps the output to the one-hot label, while the “standard (weak clamp)” model nudges the output with *β* = 1. Contrastive models use two usual equilibrium propagation phases^48^: (i) a free phase (*β* = 0 or unclamped output), and (ii) a second phase that is either weakly nudged (*β* > 0) or fully hard-clamped. Note that the contrastive (weak clamp) model is simply labelled as “contrastive” in Fig. 4. These ablations, of the number of phases and clamp type, were performed to isolate the impact of each hyperparameter. We make two main observations. First, we find that, across both datasets, the two-phase models substantially outperform the single-phase models. The standard (weak clamp) model is not shown for CIFAR-10, as it significantly underperformed compared to other models. Second, nudging seems to provide a significant benefit over full clamp on MNIST but not on CIFAR-10, where the models reach similar performance.

## 1 Properties of predictive dendrites model

In this section we describe the properties of three key variables in the PDM, which we summarize in three corresponding Lemmas. First, we show that the backward weights tend to converge during learning towards the inverse of the forward weights. Second, we show that when a trained network makes a prediction, the neural activity tends to be equal to the feed-forward input (analogously to artificial neural networks). Third, we derive the expression for prediction errors during training, and show how the errors in adjacent layers are related.

### Lemma 1.

*In a linear PDM with n neurons in each layer, if energy of all neurons reaches 0 during relaxation for a sufficiently large number of distinct patterns of neural activity (including n patterns that are not linear combinations of each other), then:*

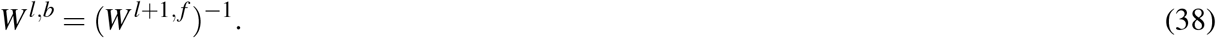

Such relationship between weights naturally arises during training, because only such backward weights enable minimisation of energy to 0.

To prove this lemma, we first note that when all prediction errors are equal to 0, then **x**^*l*^ = **v**^*l,b*^ and **x**^*l*+1^ = **v**^*l*+1, *f*^, so using the definition of backward dendritic potential **v**^*l,b*^ = *W*^*l,b*^**x**^*l*+1^, we obtain:

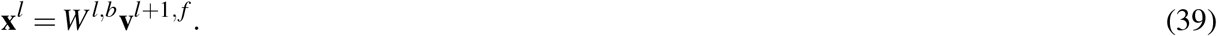

Using the definition of forward dendritic potential **v**^*l, f*^ = *W*^*l, f*^ **x**^*l*−1^, we see that the activity of neurons in layer *l* must satisfy:

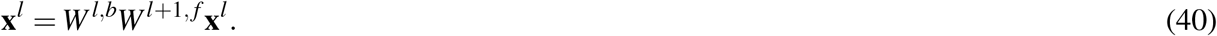

This property holds when the product *A* = *W*^*l,b*^*W*^*l*+1, *f*^ is an identity matrix. The identity *A* is a unique solution if the above equation must be satisfied for *n* patterns of neural activity **x**^*l*^ that are not linearly dependent (matrix *A* has *n*^2^ elements, so Eq 40 could be viewed as a set of *n* equations with *n*^2^ unknown; each independent pattern **x**^*l*^ for which this equation must hold provides further *n* constraints, so with *n* patterns we have *n*^2^ equations that uniquely constrain *n*^2^ elements of *A*). Matrix *A* being an identity implies Eq 38, which concludes the proof.

It is worth clarifying that the above Lemma does not make assumptions on how the distinct patterns are evoked (they may be produced e.g. by constraining both inputs and outputs during training, or by just constraining input neurons).

Supplementary Fig. 7 illustrates that during training of a sample linear PDM, the backward weights approach the inverse of forward weights.

The next two properties concern the activity of neurons during prediction and training, respectively. Hence, let us indicate with a superscript **x**^*⊖*^ that the neural activity was generated during prediction where only the input layer was constrained, and superscript **x**^*⊕*^ that the neural activity was generated during training when both the input and output layers were constrained.

**Supplementary Fig. 7.**
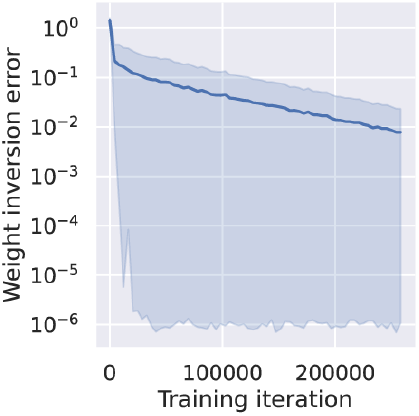
Convergence of backward weights to the inverse of forward weights during training. The weight inversion error is defined as 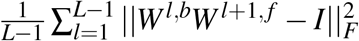, where *I* denotes an identity matrix and ||.||_*F*_ denotes the Frobenius norm. The network has 1 hidden layer with 2 neurons per layer. The weights were initialized from a uniform distribution between −1 and +1. The network was trained on inputs drawn from a uniform distribution between 0 and 1, with the size of 2 neurons, and the output is set to be the same as inputs. The experiment was repeated 10 times. Bands represent the 68% confidence intervals.

The second property is that the PDM propagates activity of neurons during prediction, as in feedforward neural networks.

### Lemma 2.

*In a linear PDM, if feedback weights are inverse of forward (Eq 38), then the activity of neurons during prediction satisfies:*

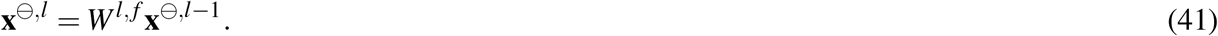

This lemma can be proven by mathematical induction over layers. In the first step of induction, we notice that Eq 41 is satisfied for *l* = *L*, as the output neurons have only single dendrite. In the second step of induction, we will show that if Eq 41 is satisfied for *l* + 1, it is also satisfied for *l*. The activity of hidden neurons in the PDM needs to satisfy:

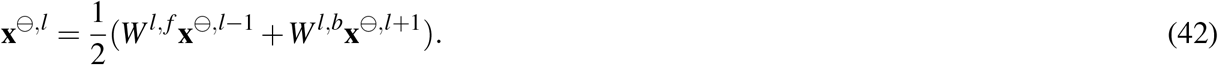

We can use Eq 41 to express **x**^*⊖,l*+1^ in Eq 42 and obtain:

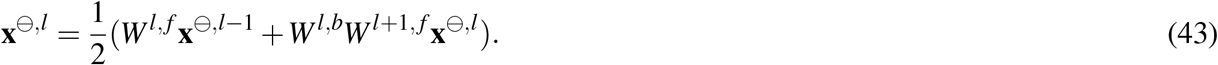

Since we assumed that the backward weights are inverses of forward (Eq 38), terms *W*^*l,b*^ and *W*^*l*+1, *f*^ cancel, and solving the above equation for **x**^*⊖,l*^ we obtain Eq 41. This concludes the proof.

The third property of the PDM concerns dendritic errors, which we define for layer *l* and forward dendrite as *ε*^*l, f*^ = **x**^*l*^ −**v**^*l, f*^. We will show that these errors are equal to the difference between the target propagated backward and the input propagated forward:

### Lemma 3.

*In a linear PDM, if the backward weights are inverses of forward weights (Eq 38), then the dendritic errors associated with forward weights satisfy:*

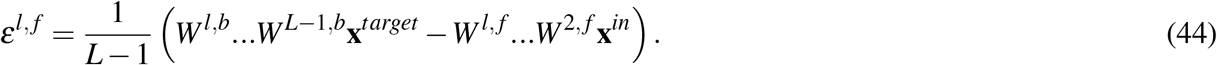

Let us first note that the activity of neurons during training must satisfy the same relationship as in prediction (Eq 42):

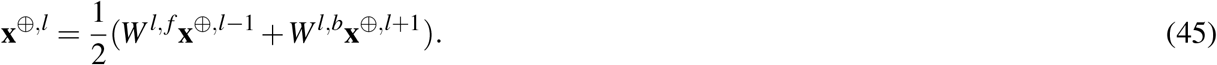

Before proving this lemma in general, let us first prove it for a special case of *L* = 3, because such proof is simpler, gives intuition for the general proof, and we consider the case of *L* = 3 in most of our simulations. From Eq 45 the activity of hidden neurons satisfies:

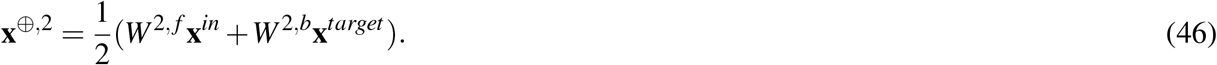

We can now evaluate the dendritic errors in the hidden layer:

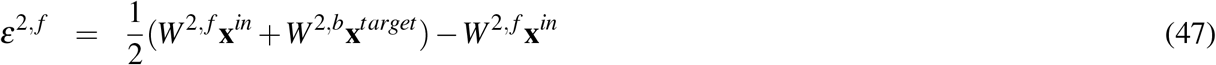

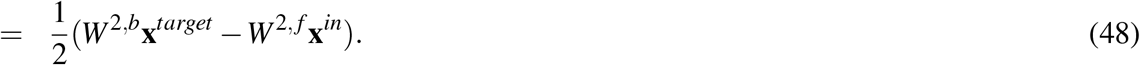

The above dendritic errors satisfy Eq 44. We can now evaluate dendritic errors in the output layer:

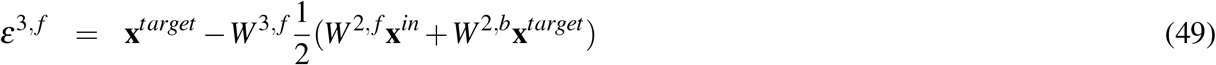

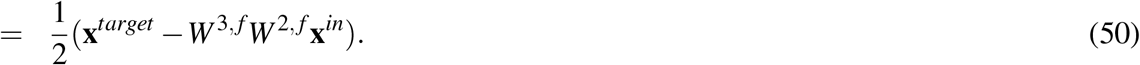

In the transition to the second line, we used Eq 38 (as it implies *W* ^3, *f*^ *W* ^2,*b*^ = *I*). The output dendritic error also satisfies Eq 44, which completes the proof for the case of *L* = 3.

A general proof for a network of any depth is analogous, but the notation becomes more complex. For a network with *L* layers, the relationship of neural activities in adjacent layers in Eq 45 results in a set of *L* − 2 equations with *L* − 2 unknowns (**x**^*⊕*,2^,…, **x**^*⊕,L*−1^). Since these equations are linear, they can be easily solved for any given *L*, and it is particularly insightful to write down the solution for *L* = 5:

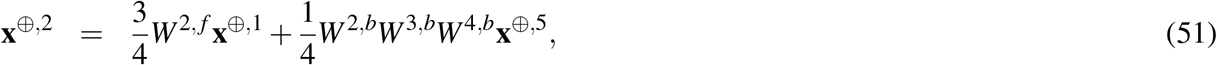

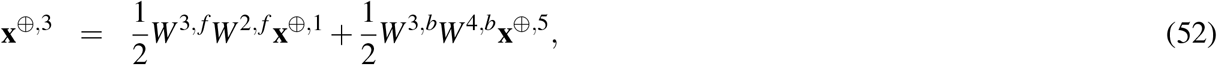

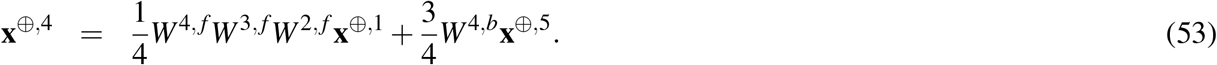

These equations illustrate that the activity of hidden neurons in an equilibrium is a weighted sum of the activities of input and output layers transformed by the weights of connections coming from these layers, and, additionally, that neurons closer to the input layer are to a larger extent influenced by the inputs, whereas neurons closer to the output layer are influenced more by the outputs. Based on a pattern seen above, we can make an ansatz that a general form of solution for any *L* will be:

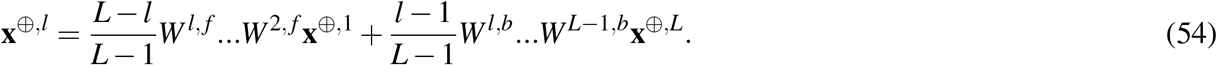

We check that this ansatz satisfies the constraint of Eq 45: Substituting **x**^*⊕,l*−1^ and **x**^*⊕,l*+1^ by expressions from Eq 54 into Eq 45, we obtain **x**^*⊕,l*^ given by Eq 54. Next, we compute the dendritic errors associated with dendrites receiving forward projections *ε*^*l, f*^ = **x**^*⊕,l*^ − *W*^*l, f*^ **x**^*⊕,l*−1^. Substituting Eq 54, we obtain Eq 44, which completes the proof.

It is worth noting that although the activity in equilibrium had different weighting of contributions of input and output layers depending on their distance (Eq 54), remarkably, these contributions are subtracted with equal weights in the dendritic errors of Eq 44. It is also interesting to note that the proportionality constant 1*/*(*L*− 1) in the expression for dendritic errors (Eq 44) depends on the number of layers. This implies that the dendritic errors are lower in the PDM with more layers, which could intuitively be explained by the fact that in the PDM the errors are redistributed in the network during relaxation. In particular, if following a prediction based on input, the output neurons are set to a target pattern, then the energy that is initially accumulated in the output layer is spread throughout all layers during relaxation.

A corollary of the Theorem is that there exists a relationship between dendritic errors in adjacent layers. Namely, the expression for dendritic errors (Eq 44) together with the requirement for backward weights being inverse of forward (Eq 38) imply the following relationship between dendritic errors in adjacent layers:

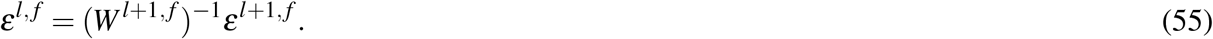

This relationship is similar to that in backpropagation, but here the errors in the next layer are multiplied by an inverse of a weight matrix (rather than its transpose as in the original backpropagation).

## 2 Target alignment of predictive dendrites model

To demonstrate formally that learning in the PDM improves its predictions, we analysed its target alignment, which quantifies how well modification of the weights changes the output of the network towards the target^20^. It is defined as a cosine of the angle between the direction in which the output changed during learning, and the direction in which it should have changed, i.e., toward the target (Supplementary Fig. 8a). Hence, if learning changes output towards target, the two directions are perfectly aligned, and target alignment is equal to 1. We show below that the absolute value of target alignment of the PDM is equal to 1 once the backward weights become the inverses of forward weights. Supplementary Fig. 8b illustrates that under these conditions, the target alignment is indeed almost always equal to −1 (there is a small probability of target alignment being equal to 1, but this occurs in the linear PDM due to a possibility of activity becoming negative - see below - which would not happen in more biologically realistic models with activity constrained to be non-negative). To investigate the robustness of target alignment to backward weights not being a perfect inverse of forward weights (as is likely to happen before learning converges), Supplementary Fig. 8c shows how the target alignment changes if noise is added to backward weights. In this case, the target alignment may decrease, but is still relatively close to 1 even for substantial deviations in the weights, suggesting that the model still modifies the weights to reduce the error on the output.

Previous work has stated a simple proof that the target alignment of target propagation is equal to 1^20^, and here we show that an analogous proof also applies to the PDM. To analyse target alignment, we will need to refer to several variables at different stages of learning, so let introduce additional notation. Let us denote the weights before update by *W* and weights after a single iteration of training on a single pair of input and target patterns (**x**^1^, **x**^*target*^) by *W*′. Additionally, let us indicate which weights were used to determine the activity of hidden and output neurons by a subscript, i.e. denote the activity resulting from weights before update by **x**_*w*_, and the activity resulting from updated weights by 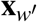. Using this notation, the updated forward weights are defined as:

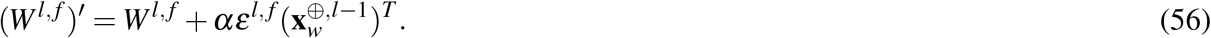

### Theorem 1.

*In a linear PDM, if before training the backward weights are equal to the inverse of forward weights (W*^*l,b*^ = (*W*^*l*+1, *f*^)^−1^*), during the training the forward weights are updated for a single iteration (Eq 56), and backward weights are updated to satisfy (*(*W*^*l,b*^)′ = ((*W*^*l*+1, *f*^)′)^−1^*), then the absolute value of target alignment is 1*.

**Supplementary Fig. 8.**
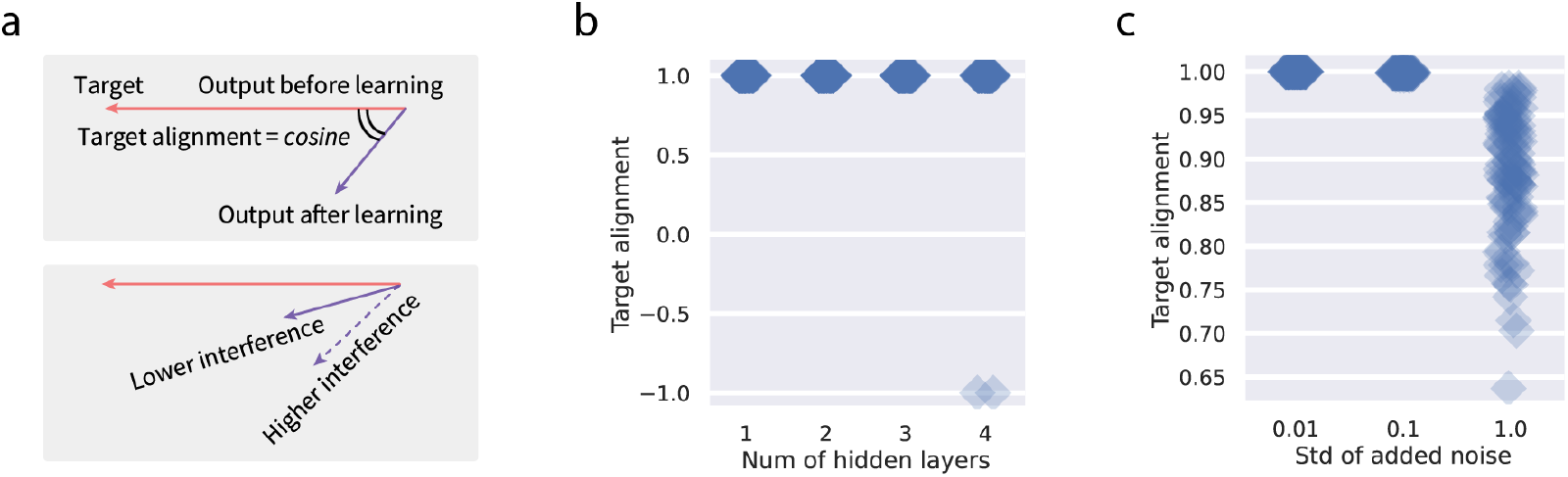
Target alignment in the PDM. ▶ **a**| Definition of target alignment. Without the loss of generality, consider a neural network with two output neurons, so that the activity of output neurons can be represented as a point on a plane. The target alignment is defined as the cosine of the angle between a vector describing the change in output due to learning 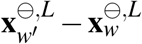, and the desired direction of change 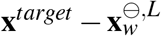. ▶ **b**| Target alignment of PDM with different numbers of hidden layers. The networks had 10 neurons per layer. The forward weights were initialized from a uniform distribution between − 0.1 and +0.1. The backward weights were initialized to be the inverse of the corresponding forward weights. The network was trained on inputs and outputs drawn from a uniform distribution between 0 and 1, and the size of both are 10 neurons. After a single iteration of training, the target alignment was computed. Each experiment are repeated 100 times, corresponding to each square in the figure. ▶ **c**| Effects of adding noise to backward weights on target alignment. The network with 1 hidden layer was trained as in panel b, and additionally random numbers from a normal distribution with mean 0 and standard deviation shown on the x-axis were added to the backward weights after they were set to the inverse of the forward weights and before actual learning taking place.

Before providing a proof of this theorem, let us comment on the assumptions it makes. It assumes that before training the backward weights are equal to the inverse of forward weights, but we explained in the previous section that the weights tend to converge to this relationship, so after a period of training they are likely to approximate it. Second, the theorem assumes that the same relation between forward and backward weights holds after training. This assumption is likely to approximately hold if the relationship was present before learning and the learning rate is small, because then the weights would not be changed much by learning. Nevertheless, we demonstrate numerically in Supplementary Fig. 8c, that the target alignment is close to 1 even if these assumptions are not fully satisfied.

To prove the theorem, let us calculate the activity of neurons in the output layer when the model makes a prediction after training. From Lemma 2 it must satisfy:

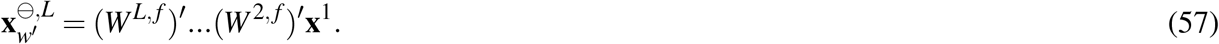

Using Eq 56, the above expression becomes:

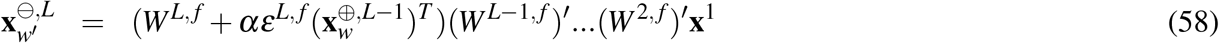

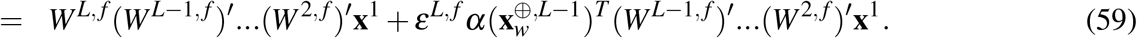

Note that 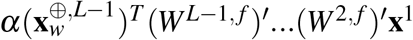 is a scalar, so let us denote it by:

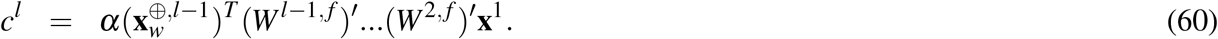

Using this notation and Eq 56 again, the network output can be written as:

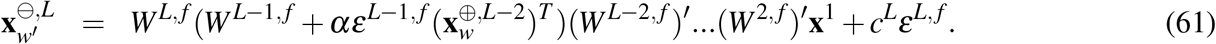

Expanding the bracket and using Eq 55 we get:

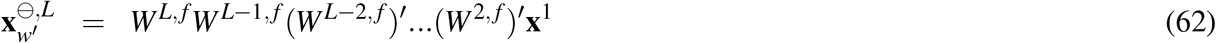

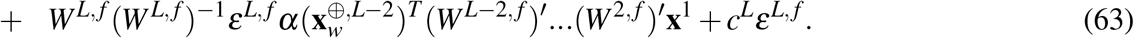

In the second line, note that *W*^*L, f*^ and (*W*^*L, f*^)^−1^ cancel, and using the notation defined in Eq 60, we obtain:

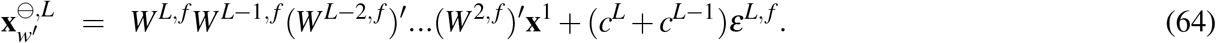

Continuing the above operations we eventually obtain:

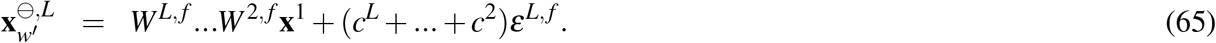

According to Lemma 2, the above expression is equal to:

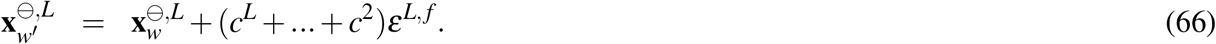

The above equation implies that the change in output 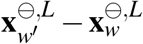 is proportional to the error in the output layer, which according to Eq 44 is equal to 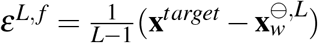, which is proportional to the direction towards target. This implies that the absolute value of target alignment is 1, which completes the proof.

The target alignment of the PDM is equal to 1 if *c*^*L*^ + … + *c*^2^ > 0, but theoretically it also can be equal to −1 if *c*^*L*^ + … + *c*^2^ < 0. Nevertheless, scalars *c*^*l*^ are more likely to be positive, because according to their definition in Eq 60, they can be written as (using Lemma 2):

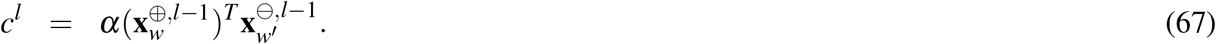

Hence *c*^*l*^ tends to be positive as long as most of the neurons do not change the sign of their activity between training and subsequent prediction. Consequently, we observed in simulations (Supplementary Fig. 8c) that the target alignment was very rarely equal to −1, and almost always equal to 1.

## 3 Relationship to bidirectional predictive coding

In this section, we analyse a relationship between the PDM and a variant of predictive coding model, called bidirectional Predictive Coding (bPC)^21^. Specifically, we identify conditions under which both models have the same equilibria of neural activity and hence modify weights in the same way. This relationship is useful because bPC achieves equally good performance as backpropagation in standard classification tasks^21^, so the identified conditions can guide how the PDM can be adjusted to achieve equally good performance.

We will first review the bPC model, then highlight its similarities and differences with PDM, and finally identify conditions under which the differences disappear.

bPC also describes a network of neurons organized into layers with activity **x**^*l*^. These neurons also receive predictions on their activity from forward (**v**^*l, f*^) and backward (**v**^*l,b*^) directions, which are defined in the same way as in PDM. bPC also defines prediction errors in the same way (*ε*^*l, f*^ = **x**^*l*^ − **v**^*l, f*^ and *ε*^*l,b*^ = **x**^*l*^ − **v**^*l,b*^). In both models, we can define an energy of a neuron as:

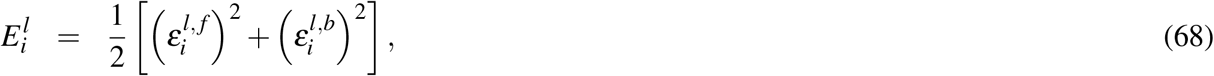

as well as the energy of the entire network:

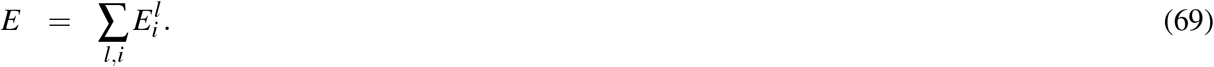

The difference between the models lies in which of the above two energies is being minimized by neural dynamics. Specifically, in the PDM, each neuron evolves its activity to minimize its corresponding energy 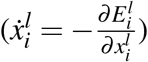, resulting in the following neural dynamics:

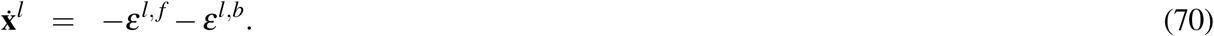

By contrast, in bPC each neuron minimizes the total energy 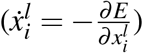, resulting in additional terms in the dynamics:

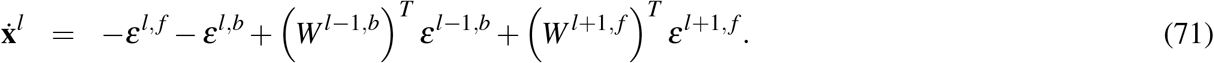

Although synaptic weight modification in each of the models minimizes its corresponding energy, the weight update rules end up being the same; hence the only difference between the models lies in dynamics of neural activity.

Comparing the above two equations highlights the key disparity between the models. Each neuron in the PDM only minimizes the errors within that neuron, while in bPC they also explicitly reduce the errors associated with other neurons in adjacent layers. This gives bPC an advantage in learning. However, we will now demonstrate that under certain conditions the activity in both models may converge to the same values.

### Theorem 2.

*If the backward weights are equal to the inverse of forward weights (W*^*l,b*^ = (*W*^*l*+1, *f*^)^−1^*), and the forward weights are orthonormal (*(*W*^*l, f*^)^*T*^*W*^*l, f*^ = *I), then the neural activity in a linear PDM and bPC converge to the same equilibria*.

Before providing a proof of this theorem, let us comment on the assumptions it makes. Analogously to Theorem 1, it assumes that the backward weights are equal to the inverse of forward weights, but as we explained earlier, the weights tend to converge to this relationship, so after a period of training they are likely to approximate it. Moreover, the theorem assumes that the weights are orthonormal, which corresponds to two requirements: (i) that the length of weight vector of each neuron is 1, and (ii) that the weight vectors are orthogonal. The first requirement may be easily enforced by a normalization that could be biologically implemented by homeostatic processes. The second requirement would be promoted by neurons having uncorrelated activity, which can be arise in biological networks through inhibitory connections among neurons^77^.

To prove this theorem, we first note that in equilibria the activity does not change 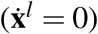, hence the neural activities in the two models need to satisfy Eq 70-71 with left hand sides set to 0, i.e., the activity in the PDM needs to satisfy:

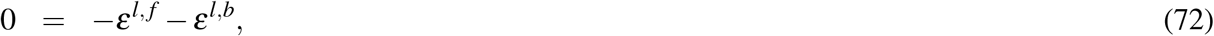

and in bPC the activity must satisfy:

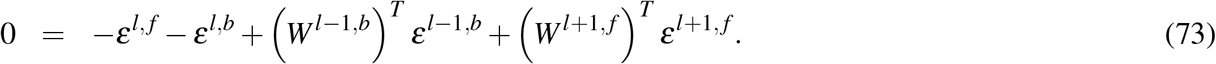

To help understand the equilibria satisfying these equations, it is helpful to consider a mechanical analogy introduced in Supplementary Fig. 1. Supplementary Fig. 9 illustrates the forces acting on a neuron within bPC in hidden layer *l* after a network converged to an equilibrium during training. There are four forces corresponding to the four terms in Eq 73, and let us define them as: *F*^*l, f*^ = −*ε*^*l, f*^, *F*^*l,b*^ = −*ε*^*l,b*^, *F*^*l*−1,*b*^ = *W*^*l*−1,*b T*^ *ε*^*l*−1,*b*^, *F*^*l*+1, *f*^ = *W*^*l*+1, *f T*^ *ε*^*l*+1, *f*^. Forces *F*^*l, f*^ and *F*^*l,b*^ come from the springs attached to the neuron, whereas forces *F*^*l*−1,*b*^ and *F*^*l*+1, *f*^ come from springs in adjacent layers (so in our mechanical analogy of bPC, neurons are connected to the adjacent layers via stiff rods). These latter forces (*F*^*l*−1,*b*^ and *F*^*l*+1, *f*^) are not present in the PDM (in its mechanical analogy, neurons could be thought as being connected via cables that just set the activity of input nodes in adjacent layers). Nevertheless, in Supplementary Fig. 9, force *F*^*l*−1,*b*^ pulls neurons in the same direction as force *F*^*l, f*^, and similarly force *F*^*l*+1, *f*^ pulls in the same direction as *F*^*l,b*^. In fact, in the case illustrated in Supplementary Fig. 9 where all weights are equal to 1, the forces are equal:

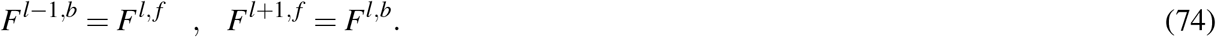

**Supplementary Fig. 9.**
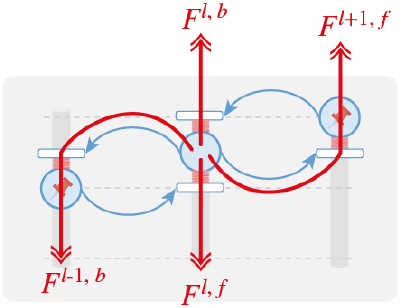
Forces acting on a hidden node in sample bPC.

Although terms *F*^*l*−1,*b*^ and *F*^*l*+1, *f*^ do not appear in the dynamics of the PDM (Eq 70), it is useful to consider their values, as they help inform how its equilibrium activity relates to the equilibrium in bPC. If Eq 74 is satisfied after activity converged, then the equilibrium of the PDM must be the same as of the bPC, because the sum of terms in Eq 73 will be just twice the sum in Eq 72, so the two equations must have the same solution. Let us now identify the conditions under which the second equality (*F*^*l*+1, *f*^ = *F*^*l,b*^) in Eq 74 is satisfied in the PDM. Using the corollary of Lemma 3 (Eq 55) we obtain:

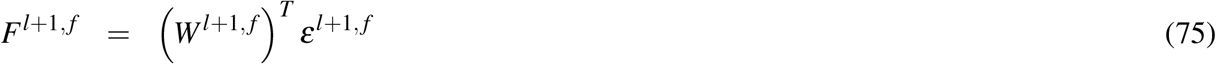

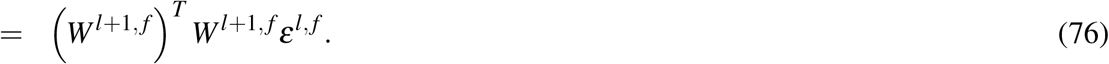

At equilibrium of PDM, the two prediction errors associated with a neuron satisfy *ε*^*l, f*^ = −*ε*^*l,b*^, hence

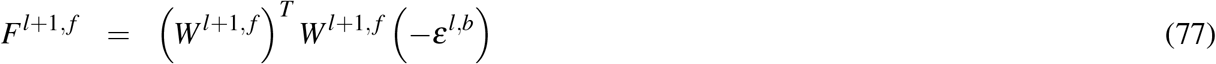

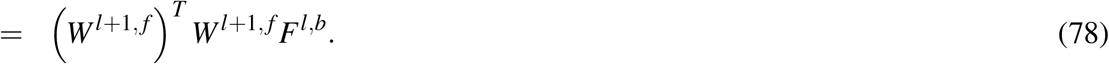

We note that if the forward weights are orthonormal, the second equality of Eq 74 is satisfied. The first equality in Eq 74 can be shown in an analogous way. This completes the proof.

## 4 Energy of a non-linear neuron

In this section, we show how the dynamics of neural activity and synaptic weights in the nonlinear PDM (shown in Fig. 3b and given by Eqs 6-8, 27) can be derived from minimisation of an energy function. For the firing-rate formulation, we consider the following augmented energy:

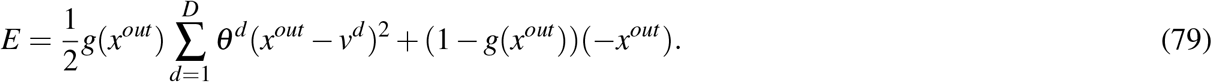

In this energy function, different dendritic errors are weighted differently by coefficients *θ*^*d*^ (such coefficients weighting dendritic errors are similar to precision parameters used in predictive coding networks^19^). The energy is also scaled by *g*(*x*^*out*^), reflecting the assumption that the prediction errors of a neuron only matter if this neuron has non-zero activity. An analogous scaling of energy or loss associated with individual neurons emerges automatically for energies or losses defined for the whole network, where changing the activity of a neuron can only reduce the global energy or loss if the activity is in a non-flat part of the neuron’s non-linear function. Therefore, we add analogous scaling to the above energy, which here is defined for a single neuron. A trivial way to minimize the first (main) term in Eq 79 is by making the neuron inactive, i.e., *x*^*out*^ = 0. Hence to prevent this, we add a second term that penalizes activity equal to zero. This second term may reflect homeostatic processes that maintain a certain level of average neuronal activity.

We assume that the neural activity evolves to reduce the energy. A negative gradient of this energy over neural activity is equal to:

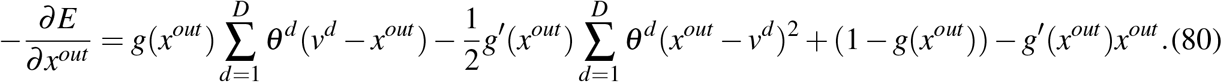

For the ReLU nonlinearity considered in this paper, *g*(*x*^*out*^) is constant except for one point where it is discontinuous, hence *g*′(*x*^*out*^) is equal to 0 except for one point where it is not defined. Therefore, we ignore the terms scaled by *g*′(*x*^*out*^) and assume that the neural activity evolves according to the resulting gradient:

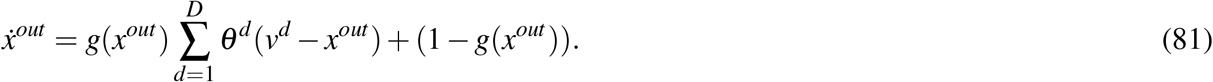

The activity to which the above equation converges depends on the sign of somatic input 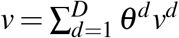. First, let us consider a case when *v* > 0. If the neural activity *x*^*out*^ is positive, then *g*(*x*^*out*^) = 1, and we can find a fixed point by noting that in an equilibrium the neural activity does not change 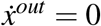:

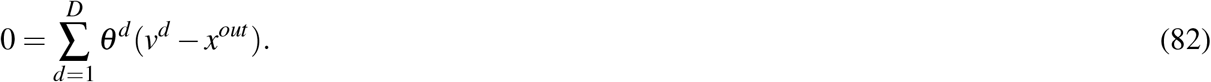

Solving for *x*^*out*^ we obtain:

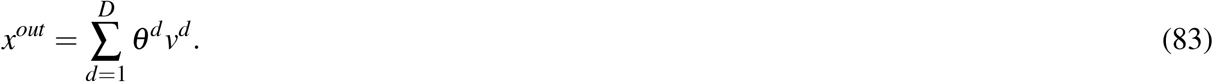

If the neural activity is equal to zero (or negative), it will increase, driven by the second term in Eq 81, until it reaches a positive value, at which point it will start converging to the equilibrium of Eq 83.

Second, let us consider a case when *v ≤* 0. If the neural activity *x*^*out*^ is positive, then it will be driven towards 0 by the first term in Eq 81. When the neural activity *x*^*out*^ reaches zero, it will be driven to a positive value by the second term in Eq 81. Thus the neural activity *x*^*out*^ will fluctuate around 0 (taking infinitesimally small values). Combining the two cases, we see that the neural activity *x*^*out*^ approaches ReLU of somatic activity *v*, as stated in Fig. 3b and Eqs 6-7.

Calculating the negative gradient of the energy with respect to a weight, we obtain:

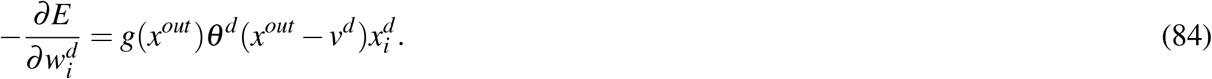

The above quantity is proportional to the weight update in Fig. 3b and Eq 8.

In the spiking formulation, the output signal is already a non-linear function of the input, because it is non-negative as the spikes can only increase the output signal above zero. Therefore, there is no need to introduce non-linearity to the energy for the purpose of deriving the activity. Nevertheless, a weight update can be obtained from a non-linear energy function, analogous to that in Eq 79, in which the energy is close to zero if the the output signal is below *γ* (we omit the differential weighting of dendrites for brevity):

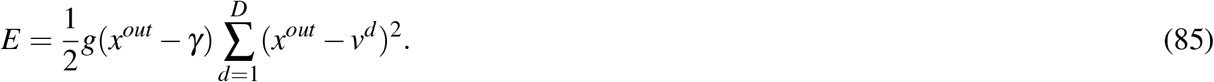

Calculating the negative gradient of the energy with respect to a weight, we observe that it is gated by the output signal, analogously as in Eq 84:

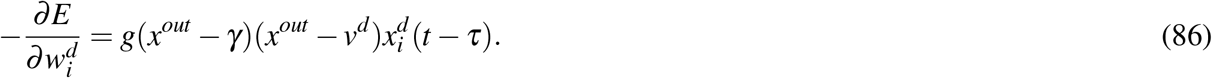

The above quantity is proportional to the weight update in Eq 27.

## 5 Representation collapse in the linear stochastic task

Below, we analyse why the weights to the hidden neuron in the network shown in Supplementary Fig. 4a collapse during learning of the simple linear stochastic task.

In this model, we explore the dynamics of the weights to the hidden neuron, *w*^2, *f*^ and *w*^2,*b*^. The model consists of three neurons (Supplementary Fig. 4a): the input neuron *x*^1^, the hidden neuron *x*^2^, and the output neuron *x*^3^. We consider the case where the activity of the hidden neuron is influenced by both the input and the output neurons equally:

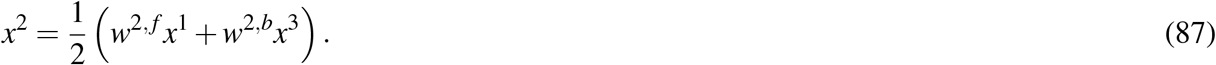

Substituting the above expression into the weight update rule from Fig. 1f, we obtain for the weight of the hidden neuron:

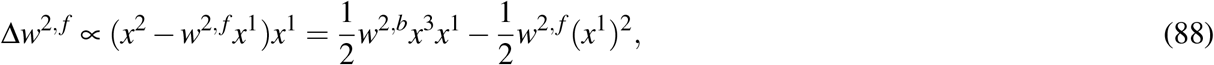

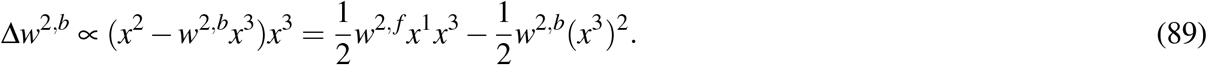

It is important to distinguish between superscript notations and power exponents. We utilize parentheses to clearly define power exponents. For example, in the expression (*x*^3^)^2^, the superscript ‘3’ indicates the third neuron, while the exponent ‘2’ outside the parentheses indicates squaring of the entire term.

We analyse the weight dynamics in a linear PDM, focusing on a scenario where the input is set to *x*^1^ = 1 and the output *x*^3^ follows a normal distribution

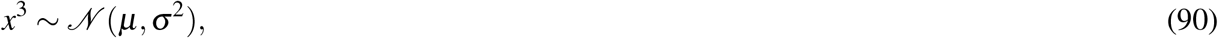

where *µ* is a non-zero constant. This setup allows us to explore how constant inputs and stochastically varying outputs influence the model’s weight adjustments.

Given that *x*^1^ = 1 and incorporating the stochastic nature of *x*^3^, we express the update equations for *w*^2, *f*^ and *w*^2,*b*^ as follows:

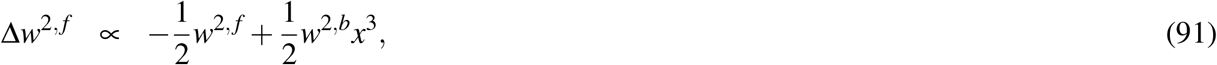

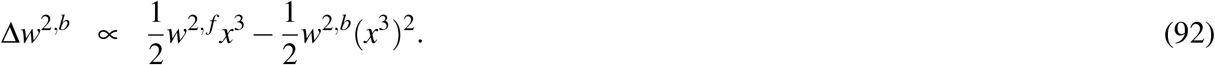

Calculating the expected values of these weight updates requires the following statistics of the output *x*^3^ during training: E[*x*^3^] = *µ*, and E[(*x*^3^)^2^] = *µ*^2^ + *σ*^2^, (found using E[*X*^2^] = (E[*X*])^2^ + Var(*X*)). The expected values of weight updates become:

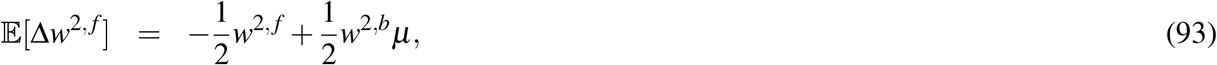

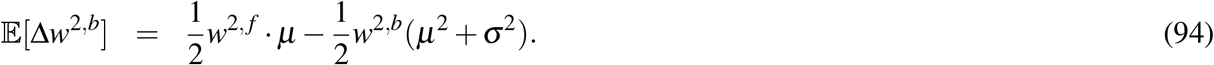

At the fixed point where the expected weight updates are zero, we derive the following system of equations:

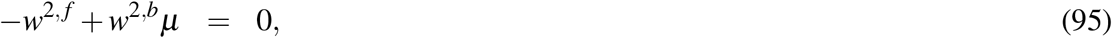

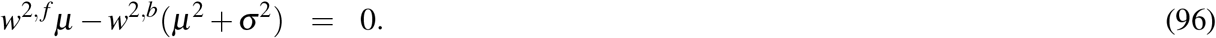

Solving these equations under different conditions for *σ*^2^ yields:

- If *σ*^2^ = 0, the system has infinitely many solutions where *w*^2, *f*^ = *µw*^2,*b*^.
- If *σ*^2^ > 0, the unique solution is *w*^2, *f*^ = *w*^2,*b*^ = 0.

This demonstrates that the presence of non-zero variance in the output noise crucially affects the stability and convergence of the weights. Specifically, any non-zero variance leads to both the forward and backward weights into the hidden neuron converging to zero, illustrating the model’s sensitivity to output noise and its significant implications for learning stability.

